# Heterogeneity in signaling pathway activity within primary and between primary and metastatic breast cancer

**DOI:** 10.1101/2020.07.27.223834

**Authors:** Márcia A. Inda, Paul van Swinderen, Anne van Brussel, Cathy B. Moelans, Wim Verhaegh, Hans van Zon, Eveline den Biezen, Jan Willem Bikker, Paul J. van Diest, Anja van de Stolpe

## Abstract

**Background:** Targeted drug treatment aims to block tumor driving signaling pathways, and is generally based on analysis of one primary tumor (PT) biopsy. Phenotypic heterogeneity within primary and between primary and metastatic lesions was investigated.

**Methods:** Activity of androgen and estrogen receptor, PI3K-FOXO, Hedgehog, TGFβ, and Wnt signaling pathways was measured in breast cancer samples using a novel mRNA-based assay platform. Macro-scale heterogeneity analysis was performed on multiple spatially distributed PT tissue blocks from 17 luminal A-like, 9 luminal B-like, and 9 ER-negative primary breast cancers; micro-scale heterogeneity analysis was performed on four “quadrant” samples of a single tissue block of respectively 9, 4, and 4 matched PT. Samples from 6 PT with matched lymph node (LN, n=23) and 9 PT with distant metastatic sites (DS, n=12) were analyzed. Statistical variance analysis was performed with linear mixed models. A “checkerboard” model was introduced to explain the observed heterogeneity in PT.

**Results:** Within PT, macro-scale heterogeneity in signaling pathway activity was similar to micro-scale heterogeneity, with a possible exception of the PI3K pathway. Variation was significantly higher on microscale for Hedgehog and TGFβ pathways. While pathway activity scores correlated significantly between different locations in the PT, positive correlations decreased between PT and LN, and even more between PT and DS metastases, including the emergence of a negative correlation for the ER pathway.

**Conclusion:** With a possible exception of the PI3K pathway, variation in signaling pathway activity within a single PT tissue block was generally representative for the whole PT, but not for DS or LN metastases. The higher variation in TGFβ and HH pathway activity on microscale suggested the presence of multiple small cancer cell clones. While analysis of multiple sub-samples of a single biopsy block may be sufficient to predict PT response to some targeted therapies, such as hormonal therapy, metastatic breast cancer treatment requires analysis of metastatic biopsies. The findings on phenotypic intra-tumor heterogeneity are compatible with currently emerging ideas on a Big Bang type of cancer evolution.

## Introduction

Cancer can be described in terms of abnormal functioning of one or more signal transduction pathways that control major cellular functions, e.g. cell division, differentiation, migration, and metabolism. Around 10-15 signal transduction pathways can drive growth and metastasis of breast cancer (1,2). In cancer, they can be activated by receptor ligands or specific DNA mutations (3–11). Signaling pathways are in principle clinically actionable, since their activity can be modified by certain drugs or other treatments. Treatment with targeted drugs is increasingly used in breast cancer, aiming to block the tumor driving pathways in the (neo)adjuvant or metastatic setting (12). However, predicting the effect of targeted therapy has generally proven to be very difficult (13). One reason is that cancer genome mutation analysis does not sufficiently predict which signaling pathways are active in an individual tumor (13,14). Further, heterogeneity in signaling pathway activity may be present within the tumor, while targeted drug choice is usually based on analysis of a single primary tumor (PT) biopsy. Similar problems exist when choosing targeted therapy for treatment of patients with metastatic disease, where therapy choice is generally based on analysis of a tissue sample from the PT, although marked genotypic and phenotypic differences between PT and distant site (DS) metastases have been described (15,16). Unfortunately, limited knowledge is available on heterogeneity in signaling pathway activity within a PT and between PT and LN and DS metastases (17,18). One reason has been lack of reliable assays to measure signaling pathway activity in formalin fixed paraffin embedded (FFPE) tissue samples used in the routine diagnostic setting.

A novel analysis method has been described to quantify signaling pathway activity in cancer. Based on Bayesian models, the method infers a pathway activity score from transcription factor target gene mRNA levels (19–22). While originally developed for use on fresh frozen tissue samples, this method was recently adapted for use on FFPE material for a number of signaling pathways, i.e. androgen (AR) and estrogen (ER) receptor, PI3K-FOXO, Hedgehog (HH), TGFβ and Wnt pathways. Using this approach, we analyzed heterogeneity in signaling pathway activity within breast PT and between PT and LN and DS metastases.

## Methods

### Patient sample series

All tissue FFPE samples have been retrospectively collected under appropriate Dutch ruling. Breast cancer molecular subtyping was performed on surgically resected PT in a standard manner by surrogate immunohistochemistry (Figure 1A,(23)). Thus, in this study the nomenclature Luminal A and B should be interpreted as Lumina A-like and Luminal B-like.

**Figure 1.**
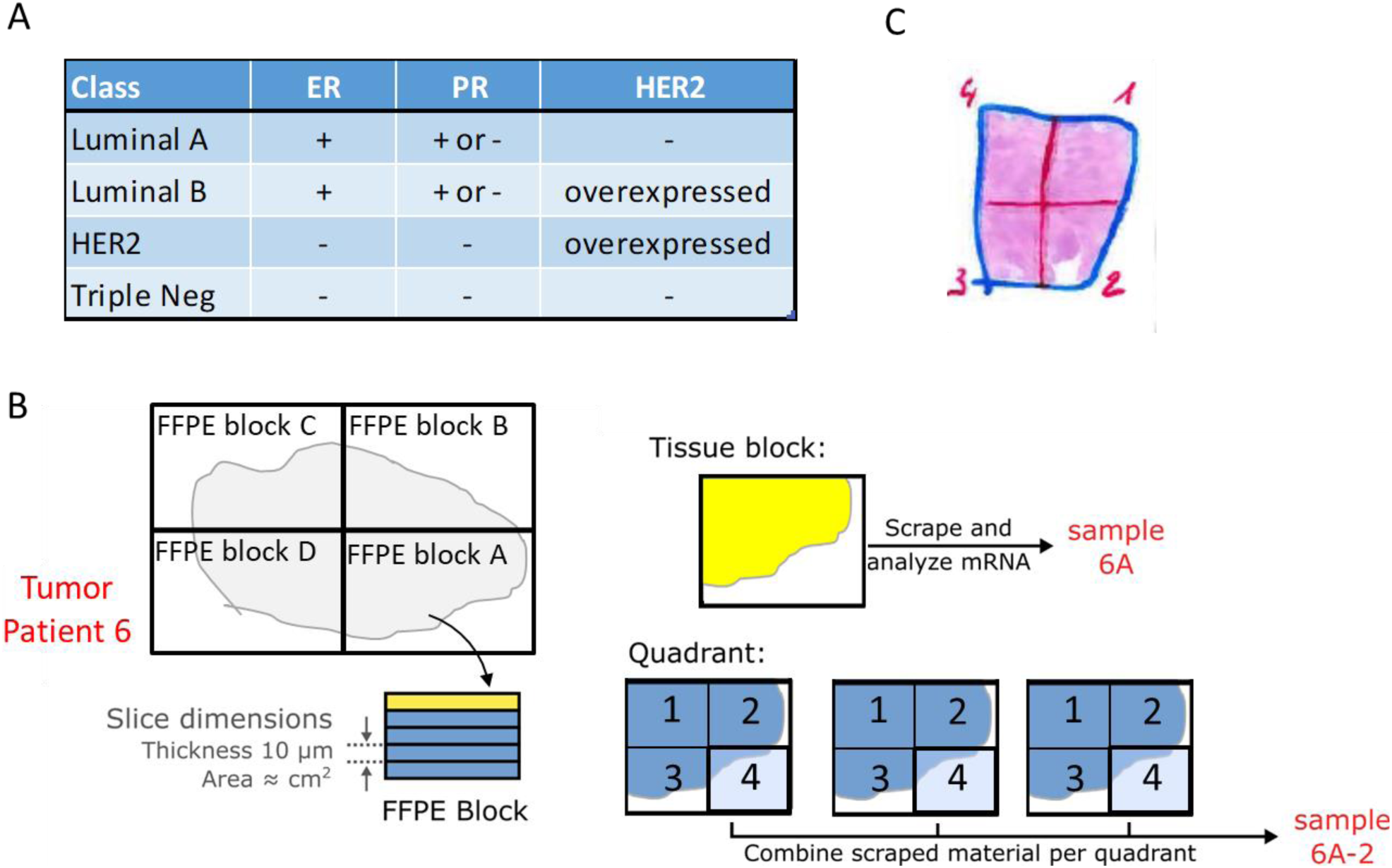
Analyzed primary breast cancer samples. A. Pathology classification of analyzed primary breast cancers. Surrogate classification of Luminal A and Luminal B subtypes based on ER/PR/HER2 immunohistochemistry staining results. HER2 overexpression in Luminal B is optional and not always present. B. Tissue samples analyzed from surgically resected primary tumors were either a tissue block sample or a quadrant sample. Tissue block samples were obtained by scrapping one or more adjacent slides (depending on the amount of cancer tissue per slide) from each available FFPE tissue block. The tumor tissue area was annotated by a pathologist and tumor-containing areas were scraped from the slide(s) for RNA isolation. Quadrant samples were obtained from one randomly selected FFPE tissue block per patient. Sequential tissue slides were divided into 4 quarts, tumor areas annotated and tissue from the quarts scraped from the slides for RNA isolation. Care was taken to scrape only cancer tissue. To obtain similar amounts of tumor tissue, multiple adjacent slides were scraped until the same area of scraped tumor had been collected. C. Typical example of a tissue block sample, showing how quadrant sample areas were divided.

### Tissue sample sets

Samples were derived from different breast cancer subtypes. A variable number of tissue blocks, taken from different locations was available per PT to investigate heterogeneity in pathway activity at the macro-scale. To investigate heterogeneity at the micro-scale, four “quadrant” samples from the same tissue block (one block per tumor) were made available for a subgroup of PT. To investigate variation in pathway activity between primary tumor and metastases, two separate patient sample sets were analyzed: one with multiple PT blocks matched with a variable number of LN metastasis samples, the other containing a single PT block and a variable number of metastasis samples from different DS. *Table* ***1*** gives a summary description of the sample sets used in this study. A detailed description with the respective sample numbers per patient is given in the Supplementary Information (Supplementary Information, SampleCounts).

**Table 1:** Summary description of sample sets used. The total number of patients and number of patients per subtype are given. Between brackets are the corresponding number of samples and their type, b: Primary tumor tissue block sample, PT: primary tumor sample, LN: lymph node metastasis sample, DS: distant site metastasis sample. Detailed description is given in the Supplementary Information (Supplementary Information, SampleCounts).

| Sample set | Description | Total | Per breast cancer subtype |  |  |  |
| --- | --- | --- | --- | --- | --- | --- |
|  |  |  | LumA | LumB | HER2 | TN |
| I | 1 to 5 block (b) samples from primary tumors | 18 (49 b) | 8 (20 b) | 5 (15 b) | — | 5 (14 b) |
| II | 2 to 5 block samples from primary tumors with 4 matched quadrant samples (per patient) | 17 (50 b) | 9 (28 b) | 4 (12 b) | 1 (2 b) | 3 (8 b) |
| III | Primary tumors (PT) and 1 to 3 matched distant metastasis samples | 10 (9 PT/13 DS) | Subtyping not available |  |  |  |
| IV | 1 to 10 lymph node metastasis from 9 patients from sample sets II and III | 9 (33 LN)* | 4 (20 LN) | 2 (3 LN) | 1 (1 LN) | 1 (8 LN) |

\* Subtyping not available from one patient.

### Sample preparation

Tissue slides were cut from the FFPE blocks at 10 μm thickness. The tumor area from which RNA was isolated was annotated by an experienced pathologist and macrodissected, aiming at similar tumor tissue volumes for RNA extraction (Figure 1B/C). RNA was isolated from the annotated areas using standard procedures, as described in detail in the Supplementary Methods.

### Measuring pathway activity

AR, ER, PI3K, HH, TGFβ and Wnt pathway activity scores were measured using biologically validated computational pathway models as described before (19–21), except that the original assays were converted from AffymetrixU133 Plus2.0 to qPCR mRNA measurements as input, to enable use of FFPE material (FIPA Pathway Plate 1.0, Philips Molecular Pathway Dx, Eindhoven, The Netherlands). For details, see Supplementary Methods and (24,25). In brief, Bayesian computational models infer the odds in favor of an active signal transduction pathway from mRNA expression levels of direct target genes of the pathway-associated transcription factor. The Bayesian computational pathway describes (i) how the expression of the target genes depends on the activation of the respective transcription complex, and (ii) how qPCR results depend in turn on the expression of the respective target genes (Figure 2). The reported pathway activity scores are presented on a 0-100 scale, where 0 corresponds to the lowest and 100 corresponds to the highest odds in favor of an active pathway that a specific model can theoretically infer. Practically this means that the actual measured range of pathway activity scores for a signaling pathway assay when performed on a specific cell type covers a reproducible part of this 0-100 scale. For the PI3K pathway, activity is inversely related to the measured FOXO transcription factor activity score, on the premise that no cellular oxidative stress is present; for this reason, the FOXO activity score is interpreted in combination with SOD2 target gene expression level to distinguish between *growth control-* and *oxidative stress*-induced FOXO activity, as described before (20,26).

**Figure 2.**
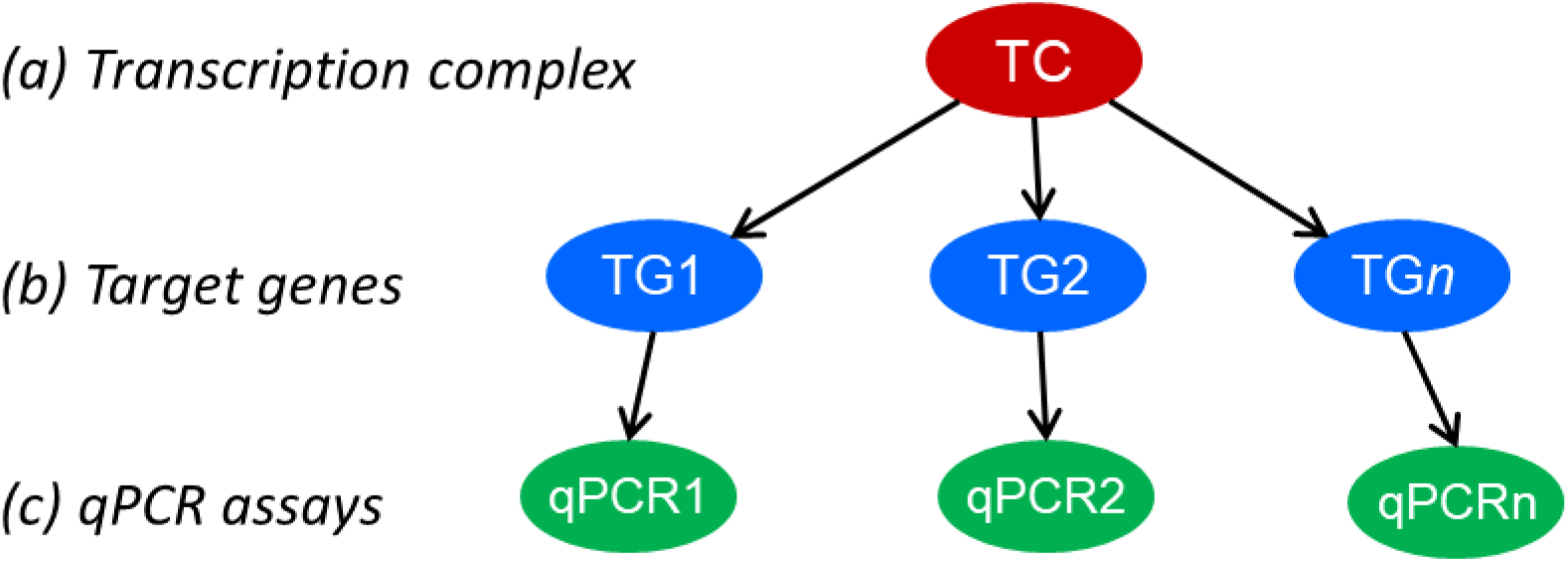
The structure of the Bayesian networks used to model the transcriptional program of signaling pathways. The transcription complex (TG) refers to the transcription factor associated with a specific signal transduction pathway, which can be present in an inactive or active gene-transcribing state; target genes (TGs) refer to direct target genes of the transcription complex; qPCR assays (PCR) refer to qPCR assay used for measuring the for the respective target gene mRNA expression level.

### Statistical data analysis

Because of the heterogeneous sample set, we used linear mixed models (27,28) to enable the best possible quantification and statistical underpinning of the pathway analysis results. In view of readability, only a brief overview of the statistical approach is given here, for an extensive description including detailed analysis results, we refer to Supplementary statistics. The model used to analyze heterogeneity in primary breast cancer subtypes considers patients grouped by cancer subtype classification, with multiple tumor block measurements per patient. The pathway activity score of a given pathway is modelled as *y* _*pq*_ = *μ* _*t*_ + *α* _*p*_ + *β* _*pq*_(Figure 3A). Here *y* _*pq*_ is the pathway activity score of patient *p* for tumor block *q*; *μ* _*t*_ the average of scores in this subtype classification group *t* (e.g. LumA or LumB); *α* _*p*_ is a random contribution of the patient tumor as a whole, compared to the group average (assumed normally distributed around zero with standard deviation *σ*_pat_); and *β* _*pq*_is the random contribution of block q which is used to accommodate tumor heterogeneity for patient p (assumed normally distributed around zero with standard deviation *σ*_block,*t*_). The value *σ*_block,*t*_ describes the spread of scores within a single patient tumor (the tumor heterogeneity) for patients in a specific breast cancer subtype *t* and *σ*_pat_ describes the variation between individual patients regardless of the breast cancer subtype they belong to.

**Figure 3.**
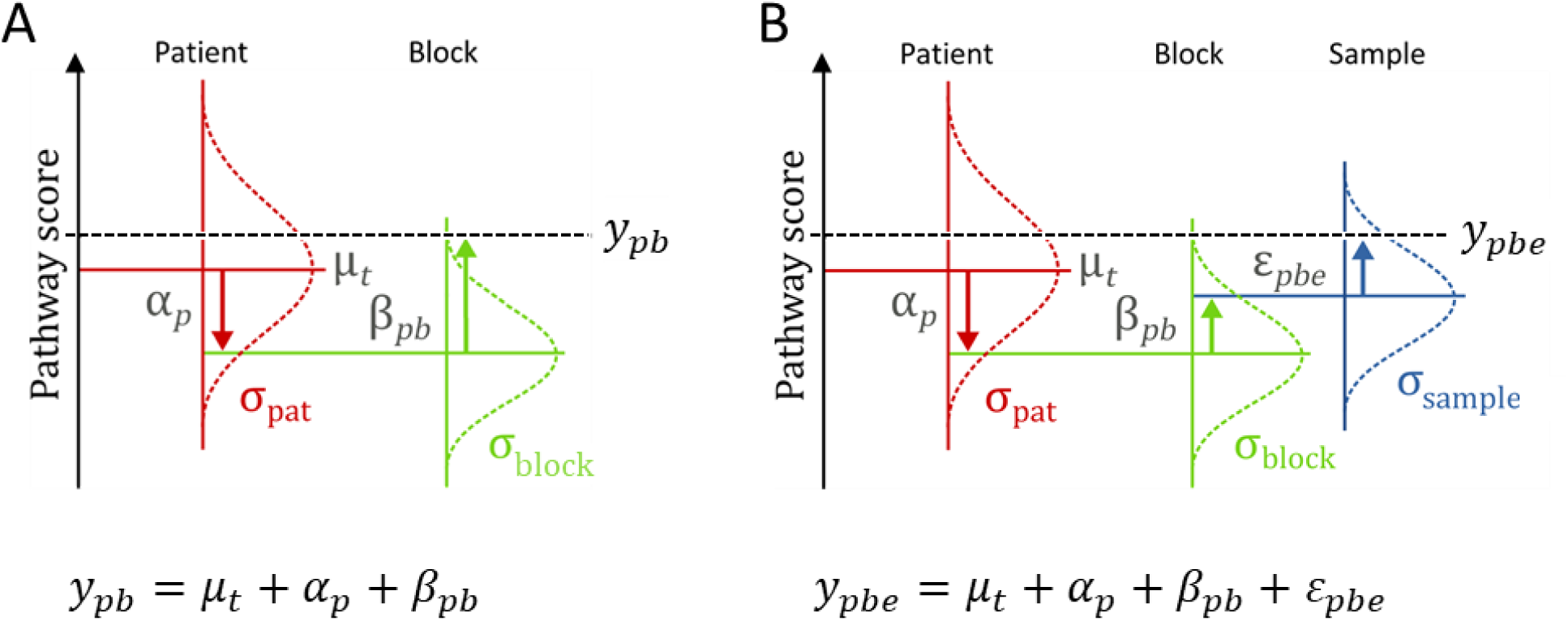
Schematic representation of models used for data analysis. A. Model used for estimation of variation in signaling pathway activity between breast cancer subtypes (y _pq_ = μ _t_ + α _p_ + β _pq_). B: Extended model enabling additional comparison between variation on micro-scale and macro-scale (y _pqe_ = μ _t_ + α _p_ + β _pq_+ε _pqe_). The models have random contributions that are added to each other; the random character of each level of contribution is emphasized by the Gaussian distribution, with indicated standard deviation (σ) and example realizations from that distribution. The models describe the pathway activity scores y _pb_ and y _pbe_ as a sum of contributions. Starting from the average score μ _t_ (computed for each breast cancer subtype t separately, with t being Luminal A-like, or Luminal B-like, or ER negative), a patient-specific contribution α _p_ (the red arrows) and a block-specific contribution β _pb_ (the green arrows) are added to the average signaling pathway activity score (μ _t_). In the extended model, an additional contribution ε _pbe_ (the blue arrow) is added to the pathway activity score of each block, to model the possibility of quadrant-to-quadrant heterogeneity. (See Figure 1B for block and quadrant definitions). A more detailed explanation is given in Supplementary statistics.

The measured pathway activity scores, in combination with this statistical model structure, are used for the statistical estimation of the parameters like *σ*_block,*t*_ or *σ*_pat_. Since they are fitted parameters to the data, the models provide a confidence interval for these quantities. For the analysis performed with quadrant measurements, the model was extended to accommodate and estimate additional micro-scale heterogeneity. The model used has an additional layer that enables taking both the tissue block and the quadrant sample measurements into account: *y* _*pqe*_=*μ* _*t*_ + *α* _*p*_ + *β* _*pq*_+ *ϵ*_*pqe*_(Figure 3B). Macro-scale heterogeneity (from blocks) and micro-scale heterogeneity (from quadrants) are defined in terms of parameters of this model. Analysis was performed in STATA (StataCorp. 2017, Release 15) and R using nlme and emmeans packages (28– 30), For details, see Supplementary statistics.

## Results

### Signal transduction pathway activity in primary breast cancer subtypes

To compare activities of the ER, AR, PI3K-FOXO, HH, TGFβeta, and Wnt signaling pathways between breast cancer subtypes, pathway activity scores measured on multiple blocks across one PT were averaged (Figure 4A, sample sets I and I, 35 patients). For an overview of all measured pathway activity scores, see Supplementary Information Data. For this comparison, pathway analysis results were available from Luminal A, B, and ER-negative (ER-) tumors.

**Figure 4.**
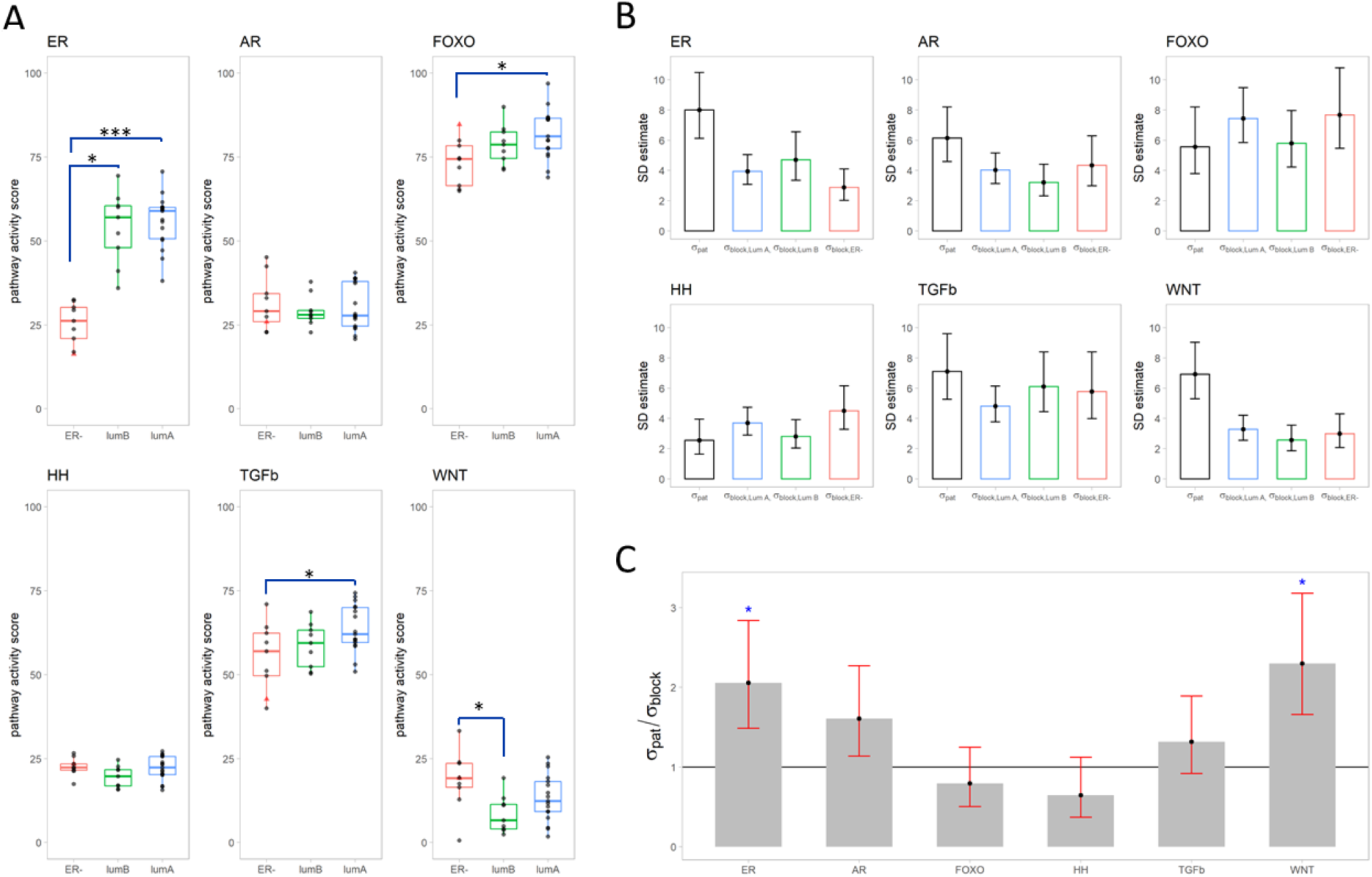
Variation in signaling pathway activity score per breast cancer subtype in the tissue block samples of sample sets I and II). A. Distribution of scores for Luminal A-like (LumA), Luminal B-like (LumB), and HER2 positive/triple negative breast cancer (ER-). Each point represents the average pathway activity score across all tissue block samples of a patient. HER2 driven patient sample depicted in red. Tukey adjusted significance levels of pairwise test for equality of means are indicated (***: p<0.0001, *: p<0.05, values in Supplementary statistics, Table S-S1). B. Estimated standard deviation (SD) with corresponding 95% confidence intervals (95% CI) of signaling pathway activity scores describing the heterogeneity in activity scores between all patients, irrespective of subtype (σ_pat_) and within a single tumor for tumors in a specific breast cancer subtype (σ_block,ER−_, σ_block,LumA_, and σ_block,LumB_), based on pathway activity scores of primary tumor tissue block samples (values in Supplementary statistics, Table S-S3). C Estimated ratios of between patient SD to within tumor SD (σ_pat_/σ_block_) with corresponding 95% confidence intervals for subtype independent σ_block_ model. Significance levels are indicated (*: p<0.05, values in Supplementary statistics, Table S-S5).

Figure 4A shows that the ER pathway had the broadest range in activity scores and largest separation in scores between subtypes. ER pathway activity scores were similar between Luminal A and Luminal B PT but markedly lower in ER-patients. FOXO pathway activity scores (as inverse readout for PI3K pathway activity) and TGFβ pathway activity scores were higher in Luminal A and progressively lower in Luminal B and ER-cancers, indicating lowest PI3K pathway activity in Luminal A tumors; while WNT pathway activity scores were higher in ER-cancers compared to Luminal type (Figure 4A, Supplementary statistics, Table S-S1).

### Variance explained per model parameter

Subsequently, we estimated the statistical variation in pathway activity between patients and within the same tumor using the model illustrated in Figures 3A and S-S1. For all signaling pathways, the variation in pathway activity within a single tumor for patients in a specific subtype (*σ*_block,t_,*t*=LumA, LumB, or ER-) was comparable for all breast cancer subtypes (Figure 4B, Supplementary statistics, Table S-S3). This implies that there is no need to compute a separate standard deviation (*σ*_block,t_) for each subtype. Instead, a simpler model, in which the estimate for the spread of scores within a single tumor is the same regardless of subtype (i.e. *σ*_block,t_ is replaced by *σ*_block_), was used in our subsequent analysis. Further analysis, using this simpler model, indicated that the variance in pathway activity score between patients (*σ*_pat_) is larger than the variance within a single tumor (*σ*_block_) for the WNT and ER pathways (i.e., *σ*_pat_/*σ*_block_ > 1), while both variances were comparable for the remaining pathways (Figure 4C, Supplementary statistics, Table S-S5).

To better understand the importance of each parameter (subtype, patient, tissue block) in the model, we also carried out a variance components analysis. Such analysis gives estimates of how much (or the percentage of) variance in activity score can be explained by each term in the model. The higher the explained percentages the higher the contribution to the total spread in the pathway activity score, the higher the importance of the parameter (details in Supplementary statistics). The results of the variance components analysis (Table 2) show that the ER pathway had the highest variation in activity score between the investigated signaling pathways, with a total variance of 284.9 (corresponding to SD=19.1). Most of that variation (78%) could be explained by the breast cancer subtype the tumor belongs to, indicating breast cancer subtype is most important parameter for the ER pathway model. This larger variance was already noticeable in Figure 4A, where the ER pathway displayed the largest range in scores. Compared to the ER pathway, total variance of the other pathways was 13 to 3 times smaller, ranging from 22.0 (SD=4.7) points on the pathway activity scale for the AR pathway to 94.4 (SD=9.7) for the TGFβ pathway (Table 2). Breast cancer subtype did not explain the observed variance in the other signaling pathways, as it did not contribute to the variation in AR pathway activity and maximally 24% for the other pathways.

**Table 2:** Variance component estimates for each signaling pathway computed using a fully nested ANOVA. Source of variance: model variable in question (indicated as Subtype, Patient, Block), Var. Comp.: variance component, variance attributed to the model variable in question. % of Total: percentage of variance attributed to the model variables taken together, StDev: variance attributed to the model variable in question, SD = √(Var. Comp.). The model variable Block indicates variation within the same tumor (σ_block_).

| Pathway | Source of variance | Var. Comp. | % of Total | SD |
| --- | --- | --- | --- | --- |
| ER | Subtype | 284.3 | 78% | 16.9 |
|  | Patient | 65.3 | 18% | 8.1 |
|  | Block | 15.5 | 4% | 3.9 |
|  | Total | 365.0 |  | 19.1 |
| AR | Subtype | 0 | 0% | 0.0 |
|  | Patient | 36.7 | 71% | 6.1 |
|  | Block | 15.0 | 29% | 3.9 |
|  | Total | 51.7 |  | 7.2 |
| FOXO | Subtype | 12.9 | 14% | 3.6 |
|  | Patient | 31.4 | 33% | 5.6 |
|  | Block | 50.1 | 53% | 7.1 |
|  | Total | 94.4 |  | 9.7 |
| HH | Subtype | 2.1 | 10% | 1.5 |
|  | Patient | 6.0 | 27% | 2.5 |
|  | Block | 14.0 | 63% | 3.7 |
|  | Total | 22.0 |  | 4.7 |
| TGFβ | Subtype | 13.1 | 14% | 3.6 |
|  | Patient | 50.8 | 54% | 7.1 |
|  | Block | 29.4 | 32% | 5.4 |
|  | Total | 93.2 |  | 9.7 |
| WNT | Subtype | 17.6 | 24% | 4.2 |
|  | Patient | 48.2 | 64% | 6.9 |
|  | Block | 9.1 | 12% | 3.0 |
|  | Total | 74.9 |  | 8.7 |

### Variation in pathway activity within a single tumor in primary breast cancer

Figure 5A illustrates range and variation of pathway activity scores measured in all tissue blocks and quadrant samples of sample sets I and II. To compare the variation of scores at macro-scale to the variation at micro-scale, pathway activity scores measured in the four quadrant samples obtained from a single FFPE block (micro-scale), as well as in all tissue blocks (macro-scale), were analysed (sample set II, 17 patients). Figure 5B depicts the spread between pathway activity scores measured in the 4 quadrant samples (micro-scale) versus the spread in scores measured in the multiple tissue blocks of the same tumor (macro-scale), and Figure 5C correlates the averaged value for pathway activity of the quadrant samples and the respective matched tissue block sample. While variation in pathway activity was observed between quadrant samples and between tissue blocks of the same tumor (Figure 5B), the averaged pathway activity scores of the quadrants strongly correlated with both the score of the matched tissue block and with the average scores of the tissue blocks (Table 3).

**Table 3.**
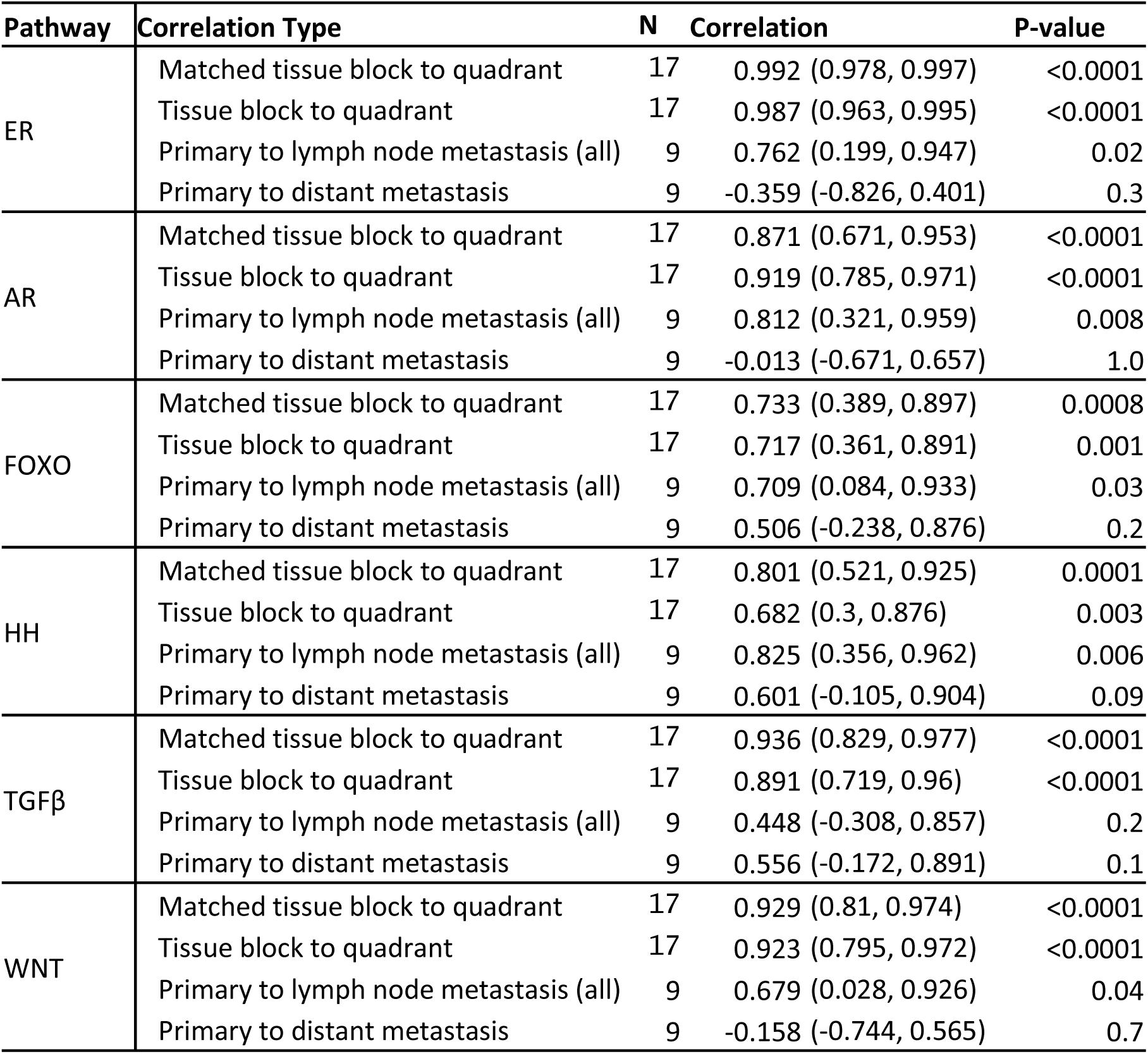
Correlation between signaling pathway activity scores for (matched) tissue block and quadrant samples (Figures 5B and 5C), for primary tumor and lymph node (Figure 6A), and for primary tumor and distant metastases (Figure 6B). For tumors with multiple samples, scores were averaged before correlation was computed. N: number of patients, Correlation given as point estimate (95% confidence interval). P-values are indicated for testing correlation=0.

**Figure 5.**
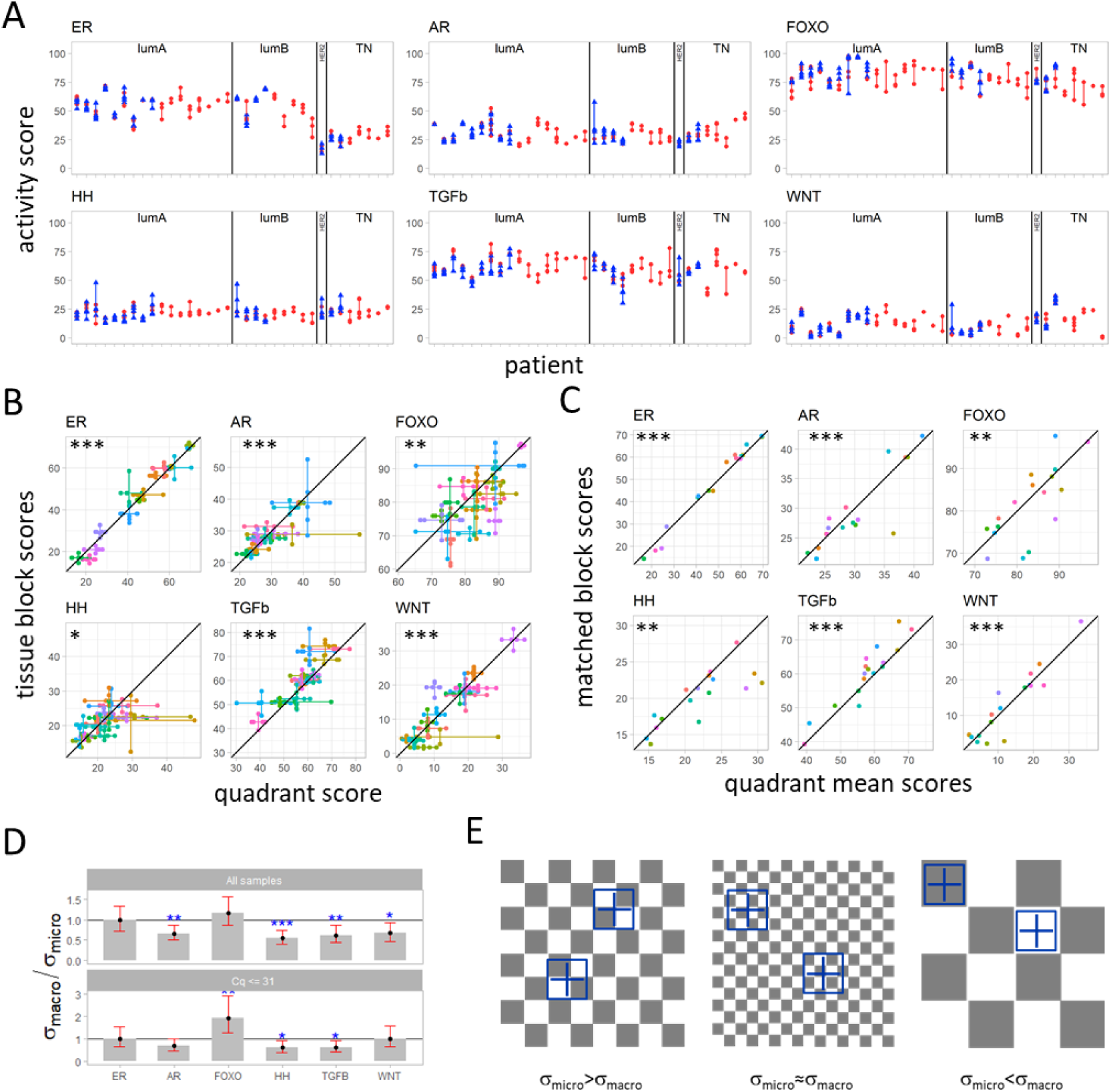
Heterogeneity in signaling pathway activity score within primary tumors at micro- and macro-scale. (Sample sets I and II) A. Pathway activity scores for tissue block (blue) and quadrant (red) samples of each patient. Ranges are presented as vertical lines; individual scores are presented as triangles. B. Spread in signaling pathway activity scores from quadrant samples (horizontal lines) vs. spread in scores from tissue block samples of the same patient (vertical lines). Significance level for the correlation between mean scores of tissue block samples versus mean scores of quadrant samples (point where horizontal and vertical lines cross) are indicated in the figure by stars (*: p<0.05, **: p<0.01, ***: p<0.001, corresponding values are in Table 3). C. Correlation between mean quadrant sample pathway activity scores and score of the corresponding tissue block sample. Stars indicate the significance level of the correlations (corresponding are p values are in Table 3). (B-C) Diagonal black lines illustrate a one to one correlation. Samples of the same patient have the same color. Black lines illustrate a one to one correlation. D. Ratio between macro-scale and micro-scale standard deviation (SD) of signaling pathway activity scores (σ_macro_/σ_micro_, measured in tissue block samples and quadrant samples, respectively). Variances computed using the macro vs. micro scale model (Figure 3B, Supplementary statistics, Figure S-S2) using all samples (top) or living out all samples with average Cq values > 31 for the reference genes used for the qPCR measurements. Significance level of Wald test p-value for comparing the ratio σ_macro_/σ_micro_ to 1 are indicated by stars (p values for model run using all samples in Supplementary Statistics, Table S-S7). “Checkerboard” visualization of a primary cancer, explaining differences in heterogeneity between micro-scale and macro-scale measurements of pathway activity. Left: Squares represent (small) cancer cell clones with variable pathway activity scores, simulated by grey (high pathway activity) and white (low pathway activity). For example, analyzing HH pathway activity in a randomly localized “tissue” sample (analogous to the tissue block samples), results in a smoothed averaged pathway activity, due to canceling out of variations in pathway activity that are present in areas smaller than the sampled area. On the other hand, when taking four quadrant samples, the varying pathway activity scores in the quadrants are measured, resulting in a higher measured pathway activity heterogeneity at micro-scale. Right: Quadrangles represent large cancer cell clones, with variations in pathway activity scores. In this case it is expected that more heterogeneity will be found on the macro-scale, since the quadrants are more likely to have the same pathway activity. This might be the case for the PI3K-FOXO pathway. For detailed information on the associated statistical model, see Supplementary statistics.

To determine whether pathway activity varied more at macro-scale (tissue blocks) or at micro-scale (block quadrants), we calculated the ratio between the standard deviation of pathway activity score at macro-scale and the standard deviation of pathway activity score at micro-scale for each signaling pathway. The standard deviations were estimated using the model illustrated in Figure 3B (details in Supplementary statistics). For the AR, Hedgehog, TGFβ and Wnt pathways, variation in pathway activity score was significantly higher at micro-scale than at macro-scale, with highest statistical significance for the Hedgehog pathway, p<0.001 (Figure 5D, Supplementary statistics, Table S-S7). For the ER and FOXO pathways, the variation in score at macro-scale was comparable to micro-scale.

Extensive noise analysis was performed to identify technical noise, which might have biased the pathway activity scores, and therefore the ratio between variation at macro-scale and micro-scale (Supplementary Methods, technical noise analysis, Figures S-M2 to S-M4). High qPCR Cq values, which were more frequent in measurements of very small samples may be associated with increased technical noise. Removal of such samples resulted in loss of significance of the higher variation in AR and Wnt pathway activity at micro-scale, but did not change the results for the HH and TGFβ pathways (Figure 5D bottom). Taking this into account, we can safely draw an overall conclusion that heterogeneity in signaling pathway activity was not larger at the macro-scale (blocks) than at the micro-scale (quadrants), except for the PI3K-FOXO pathway for which a higher variation at macro-scale could not be excluded, based on the 95% confidence interval computed when the high qPCR Cq values had been removed (Figure 5D bottom). Variation in ER pathway activity was similar at micro-scale and macro-scale and variation in HH and TGFβ pathway activity was higher at micro-scale.

### Interpreting the variation in signaling pathway activity at micro-scale versus macro-scale

Since the observed higher variation in Hedgehog and TGFβ pathway activity on micro-scale compared to macro-scale seemed counter-intuitive, a hypothetical computational *checkerboard* model was developed to help explain observed results (Figure 5E, Supplementary statistics). In this checkerboard model, squares represent cancer cell clones with variable pathway activity scores, depicted in black and white, that are present across a tumor. In case cancer clones are smaller than the tissue block and around the size of the quadrant samples (Figure 5E, left), the pathway activity scores measured in (all) the tissue block samples will average out the varying pathway activity scores measured in the quadrant samples, causing the variance between tissue blocks to be smaller than the variance between quadrants. This is likely to be the case for the Hedgehog and TGFβ pathways for which the activity scores were found to dominantly vary between the quadrants of a block. Variation in ER pathway activity was relatively small and more or less similar across the whole PT. This can be explained by either a homogeneous tumor, or variations in ER pathway activity at a very small scale (very small “clones” with varying ER pathway activity; Figure 5E middle), across the whole tumor. In this case, the quadrant samples are larger than the clones with varying ER pathway activity and the differences in score between quadrants and between tissue block samples are similar. Finally, clones that are larger than the sampled blocks and quadrants, but much smaller than the whole tumor (Figure 5E, right) could explain the potentially higher variation at macro-scale in PI3K-FOXO pathway activity. A detailed explanation is provided in Supplementary statistics.

### Differences in pathway activity between primary tumors matched and lymph node metastases

To examine the question whether pathway activity in LN metastasis is similar to pathway activity in PT, multiple PT and LN matched samples per patient were analyzed (9 patients, PT tissue blocks from a subset of sample sets II and III, LN samples from sample set IV). For all signaling pathways, activity scores within the PT were positively correlated with pathway activity scores in corresponding LN metastases (Figure 6A). However, correlations were much smaller than the correlations obtained when comparing PT block and matched quadrant samples, ranging from 0.448, 95%CI (−0.308, 0.857), p=0.2, for the TGFβ pathway to 0.825, 95%CI (0.356, 0.962), p=0.006, for the HH pathway (Table 3). This indicates that pathway activity measured in PT may not reflect well the pathway activity measured in the corresponding LN metastasis.

**Figure 6.**
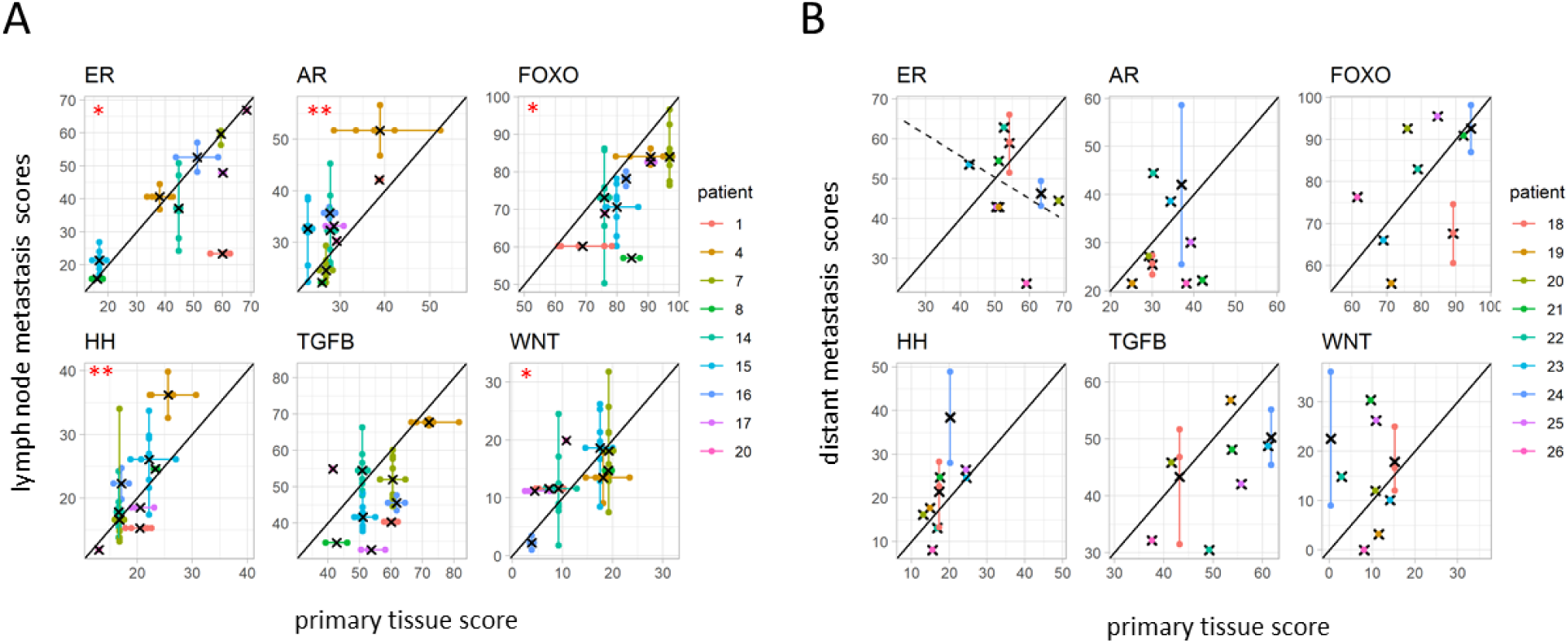
Heterogeneity in signaling pathway activity between primary tumor and metastases. A. Correlation between measured pathway activity scores in primary (x-axis) tumor and matched lymph node metastases (y-axis) visualized by plotting pathway activity scores (Sample sets IV and II/III). B. Correlation between measured pathway activity score in primary tumor (single sample) and matched distant metastases (Sample set III). Per plot, colors indicate samples that belong to one patient (patient IDs indicated in the legends), the cross indicates the average, and the black line is a visual guide for a one-to-one correlation. Significance level for the correlation between mean scores are indicated by stars (*: p<0.05, **: p<0.01, corresponding values are in Table 3).

Quantitative differences in pathway activity score between LN metastatic samples and PT samples were calculated by comparing, for each patient *p* with matched PT and LN metastatic samples, (a) the delta in pathway activity score between each *lymph node metastasis* and the average score of the respective PT samples, Δ _*p,LN*_, and (b) the delta in score between each PT tissue block and the respective PT block averages, Δ _*p,block*_ (Supplementary statistics, Figure S-S6). For all pathways, the spread in Δ _*p,LN*_ was larger than the spread in Δ _*p,block*_, indicating that difference between PT and LN metastases was larger than the variation in pathway activity within the PT (Supplementary statistics, Tables S-S8, S-S9). For the TGFβ pathway and FOXO, activity scores were generally lower in lymph node metastases compared to the PT, with lower FOXO indicated higher PI3K pathway activity; for AR, HH and WNT pathways, activity scores were more frequently higher in lymph node metastases compared to the PT (Figure 6A, Supplementary statistics, Tables S-S8, S-S9).

### Differences in pathway activity between primary tumors and matched distant metastases

For this analysis, a single PT sample (per patient, n=9) matched with a variable number of DS metastatic samples was available (sample group III). Except for the TGFβ pathway, correlations between DS metastasis and matched PT were worse than correlations between LN metastasis and matched PT. Correlations remained positive for FOXO and the HH and TGFβ pathways, but became negligible for the AR and WNT pathways, and negative for the ER pathway, ranging from positive correlation of 0.601, 95% CI (−0.105, 0.904), p=0.09 for the HH pathway to a negative correlation of -0.359, 95%CI (−0.826, 0.401), p=0.3, for the ER pathway (Figure 6B, Table 3).

To further examine the differences in pathway activity score between distant metastases and PTs, for each patient *q* with matched PT and DS metastatic samples, deltas in activity score between each *DS metastatic sample* and the only available matched PT sample, Δ _*q,meta*_, were compared to the previously computed *PT tissue block deltas*, Δ _*p,block*_ (Supplementary statistics, Figure S-S7, Table S-S9, the indices *q* and *p* emphasize the use of two different patient cohorts). Here again, the spread in Δ _*q,meta*_ was larger than the spread in Δ _*p,block*_ for all pathways, indicating that, as with LN metastasis, differences in pathway activity scores between PT and DS metastases are larger than the variation within the PT (Supplementary statistics, Tables S-S8, S-S9). While a loss of AR, ER and TGFβ pathway activity was frequently observed in metastases compared with the PT, WNT pathway activity was regularly increased in metastases (Figure 6B, Supplementary statistics, Tables S-S8, S-S9).

### Comparing pathway activity between distant metastases

Variation between distant metastases and lymph node metastases were analyzed by comparing the spread in Δ _*q,meta*_ to the spread in Δ _*p,LN*_. Apart from the TGFβ pathway, variation between distant metastases (spread in Δ _*q*,meta_) was larger than variation between lymph nodes (the spread in Δ _*p,LN*_) (Supplementary statistics, Tables S-S8, S-S9).

### Pathway activity related to metastatic organ site

Specific tissues are thought to provide a favorable niche for metastatic growth that is driven by a signaling pathway for which the activating ligand is provided by the tissue niche (31). The small number of metastatic samples from a similar organ site precluded investigation of a relation between signaling pathway activity and metastatic location. Given this limitation, pathway activity scores in individual metastatic tumors may still provide interesting information. Wnt pathway activity score was highest in bone (n=2) and ileum (n=1); Hedgehog pathway activity was highest in ovarian, ER pathway activity highest in a brain, and TGFβ highest in a skin and a bone metastatic lesion (see Supplementary Information Data).

### PI3K-FOXO pathway analysis

For only three patients, all triple negative subtype, FOXO activity was high in combination with elevated SOD2 expression level, indicating oxidative stress-associated FOXO activity, precluding direct inference of PI3K pathway activity (Supplementary methods, Figure S-M5, (20)). For all other analyzed samples PI3K pathway activity could be directly (inversely) inferred from the respective FOXO activity score.

## Discussion

To investigate variation in tumor driving signaling pathway activity within primary breast cancer tumors and between matched primary tumor and LN and DS metastases, activity of the PI3K growth factor pathway, the ER and AR hormonal pathways, and the developmental Wnt, Hedgehog and TGFβ pathways were measured at multiple locations in PT as well as in metastatic samples. Pathway activity scores were quantified using a novel method for signaling pathway analysis, adapted for FFPE samples (19–21). While initial semi-quantitative analysis of the various groups of pathway measurement results already indicated our major findings (32), the complex relationships between the available sample sets required an innovative statistical model-based data analysis approach to objectively quantify our results (28).

### Variation in pathway activity between breast cancer subtypes

As expected based on previous work, ER pathway activity was mostly defined by the breast cancer subtype and scores were much higher in PT of Luminal A and B patients, compared to ER-tumors (19,24). More interesting is that within Luminal A and B subtypes, activity of the ER pathway showed a large dynamic range. We previously showed that ER expression is a prerequisite, but not sufficient, for functional activity of the ER pathway, and the current results further confirm this (19). In the absence of activating mutations in the ER gene, the presence of ER ligands and specific downstream proteins (e.g. cofactors) determine actual activity of the ER transcription factor. This creates the necessity to measure the functional activity of the ER pathway to optimally predict sensitivity to hormonal therapy (24,33). The highest activity scores of the FOXO transcription factor, which is associated with an inactive PI3K pathway, was found in Luminal A cancers. This is in agreement with our earlier findings and with the known tumor-driving role of the ER pathway, in the absence of PI3K pathway activity, in this subtype (20).

### Within tumor heterogeneity of signaling pathway activity

To the best of our knowledge, to date no information is available on phenotypic heterogeneity within primary breast cancer with respect to functional signaling pathway activity. Knowledge on signaling pathway heterogeneity is limited to studies on variations in ER and HER2 IHC staining, and on spatial variations observed in specific gene mutations and copy number changes across the tumor (17,18,34–37). However, this does not provide information on variations in functional signaling pathway activity. In the current study, with a possible exception of the PI3K pathway, such pathway activity variation was found be comparabe both at macro-scale, across the primary tumor, and at micro-scale, within a single tumor tissue block.

The Hedgehog and TGFβ pathway showed consistently higher variation in activity at the micro-scale. Both pathways are typically active in stem cells, suggesting that multiple small clones may be present and spread throughout the PT and composed of cancer stem cells. This fits well with the cancer stem cell hypothesis, which assumes that small groups of relatively quiescent cancer cells that have acquired stem cell characteristics are present in tumor tissue (1,38,39). Both pathways are generally thought to play an important role in governing metastatic behavior and therapy resistance of cancer stem cells, another hallmark of cancer (1). Previous pathway analysis, using the same pathway activity measurement approach, suggested that overt HH pathway activity, measured in a standard tissue slide, was associated with a worse prognosis (40). In the current study, no outcome data were available, preventing further investigation of the prognostic role of activity of this signaling pathway at the micro-scale.

For the ER pathway, variation in pathway activity within the PT was lower than for the other pathways, and similar at micro-scale versus macro-scale. The variation within the primary tumor was also relatively small compared to the variation in ER pathway activity between individual luminal A/B patients. These observations can be readily explained by considering the ER pathway as the prime tumor-driving pathway in luminal breast cancer, in contrast to HER2 and triple negative cancer subtypes. Variations in ER immunohistochemistry (IHC) staining within an ER positive primary tumor have been reported, but not related to pathway activity (35). ER-activating estrogens are expected to distribute relatively evenly in tissue, which is compatible with only small differences in pathway activity as we observed before (34,41,42).

### A “checkerboard” cancer model and “Big Bang” type of cancer evolution

Our observation that, with the possible exception of the PI3K-FOXO pathway, the variation in pathway activity was generally not larger at macro-scale than micro-scale, and for the HH and TGFβ pathways even higher at the micro-scale, is compatible with currently emerging ideas on a Big Bang type of cancer evolution based on genomic analysis of PT, in which macro-scale heterogeneity appeared to be not the dominant form (17). Compatible with this model, the presented “checkerboard” model for cancer heterogeneity can be used to explain our findings in terms of variable cancer cell clone sizes, all spread more or less evenly across the tumor: the smallest “clones” or cancer cell areas with slightly varying ER pathway activity; small clones that have an active Hedgehog and/or TGFβ pathway, and possibly intermediate/large clones with varying PI3K pathway activity. In such a model, combined ER and PI3K pathway active clones may be more rapidly proliferating, while Hedgehog and TGFβ pathway active clones are “neutral”, less proliferative, and may have a stem cell character.

### Variation in pathway activity between primary and lymph node metastases

Subsequently we investigated whether pathway analysis of the primary tumor could predict pathway activity in metastatic tumors. While pathway activity scores between quadrant samples of a single block, and between blocks of one PT showed highly significant correlations, correlations decreased between PT and LN metastases and significance was lost. The lowest correlation was found between PT and DS metastases. These results provide interesting new information about breast cancer metastasis. Since LN metastatic samples were spatiotemporally close to the PT, a closer relation to the PT was expected compared to the DS metastases that are spatiotemporally further away. The PI3K pathway is generally thought to be an important metastasis driving pathway in addition to its general role as a tumor “survival” pathway, explaining the relative strong correlation in activity between PT and LN metastases (43). The observed higher variation in pathway activity between DS metastases compared to LN metastases, is probably due to LN representing a singular type of metastatic niche, while DS metastases grow in all kinds of organ niches with their respective variations in microenvironment, including ligand availability for the various signaling pathways (44,45). As a few tentative examples to illustrate this, Wnt ligand availability may have caused the one intestinal metastatic tumor to have the highest Wnt pathway activity score, while local Sonic Hedgehog ligand availability may have induced the high Hedgehog pathway activity score in the ovarian metastatic lesion (31,46,47). For the ER pathway, such heterogeneity has also been demonstrated in ER positive breast cancer patients (48).

Interestingly, a negative correlation was present for ER pathway activity between PT and DS metastases. Since only ER-positive luminal patients were included in this part of the study, a number of patients probably had received adjuvant hormonal therapy, potentially resulting in selection for ER pathway-inactive metastases (49). Unfortunately, lack of treatment information precluded further analysis of this relationship between treatment and ER pathway activity.

Obvious limitations of the study were the relatively limited patient and sample numbers, and lack of clinical treatment and outcome data. Despite this, to the best of our knowledge the study is unique in allowing a quite detailed comparison between signaling pathway activity scores within a PT and between PT and metastases.

The clinically relevant conclusions that can be drawn suggest that heterogeneity with respect to pathway activity within a single biopsy may generally be representative for the whole tumor, with the possible exception of the PI3K pathway, while variation in ER pathway activity is relatively small within the PT compared to that seen between luminal breast cancers from different patients. We cannot exclude that for the PI3K pathway, multiple spatially separated biopsies may be required to get a sufficiently complete picture on this pathway’s activity pattern across the tumor.

Pathway activity scores found in the PT predicted activity in LN metastases to some extent, especially with respect to the PI3K-FOXO pathway. Prediction of pathway activity in DS metastases was not realistic any more, also in view of the large variation between metastases in different organ locations. If possible, taking biopsies from multiple metastatic locations may therefore be preferred when considering targeted therapy for metastatic disease, to choose the most effective (targeted) drug or drug combination.

In view of its high clinical relevance, especially with respect to targeted treatment in (neo)adjuvant and metastatic setting, the current findings should be confirmed in subsequent clinical studies. We recommend that a confirmation study should: (1) optimize standardization of tissue sampling; (2) note size of the tumor and location of the sample taken in the 3D space of the tumor, to reduce the influence of variable microenvironment, e.g. oxygen concentration, immune cell infiltrate, etc.; (3) use similar amounts of cancer cells to extract RNA from. Importantly, application and further development of the statistical models that we introduced should help to improve quantification of cancer heterogeneity, necessary to objectively compare clinical studies and bring the field forward.

We believe that such studies will further complement knowledge on tumor evolution and the resulting tumor heterogeneity, enabling improvement in choosing the most effective therapy for patients with breast cancer.

## Supporting information

Supplementary information samplecounts

Supplementary statistics

## Acknowledgement

We wish to thank Paul van de Wiel for the very useful comments and suggestions.

## Funding

none

## Conflict of Interest statement

Márcia A. Inda, Anne van Brussel, Wim Verhaegh, Hans van Zon, Eveline den Biezen, and Anja van de Stolpe are employees of Philips

## Notes

### Competing Interest Statement

Marcia A. Inda, Paul van Swinderen, Anne van Brussel, Wim Verhaegh, Hans van Zon, Eveline den Biezen, and Anja van de Stolpe are (former) employees of Philips

