## Supplementary methods for "Heterogeneity in signaling pathway activity within primary and between primary and metastatic breast cancer"

### SUPPLEMENTARY METHODS (S-M)

#### RNA isolation and qPCR measurements

The RNA isolation was done at UMC Utrecht. Three consecutive coupes were cut from each FFPE block. The slides were xylene treated and the annotated (by a pathologist) tumor area was manually scraped. RNA was isolated with the Qiagen miRNeasy FFPE kit (Cat. No. 217504). Samples were shipped from UMCU to Philips Research Eindhoven, and subsequently treated with DNAse according to the protocol of the Siemens VERSANT Manual FFPE Nucleic Acid Purification Protocol, to obtain pure RNA samples. The concentration of the RNA was measured using Nanodrop. At least 100 ng RNA was used in the Philips Research Multi-Pathway RT-qPCR plate v0.1 (MPP), containing reagents for standardized performance of 96 qPCRs. The Invitrogen SuperScript® III Platinum® one-step qRT-PCR system (ref 11732-088) was used for all samples. The MPP v0.1 RT-qPCR assays had been previously validated with the addition of a PCR additive in the reaction mix. All RT-qPCR reactions were executed using the BIO-RAD CFX96 Touch™ Real-Time PCR Detection System. Each qPCR requires at least 1 ng of RNA.

#### Development of the FIPA Pathway Plate 1.0 (Philips Molecular Pathway Dx, Eindhoven, The Netherlands)

The qPCR FIPA Pathway Plate 1.0 was developed for use on formalin fixed paraffin embedded material and was calibrated and biologically validated on samples with a known pathway activity status. In the qPCR-based pathway analysis tests, the original number of measured target genes was reduced and the Bayesian interpretation model was adapted to the reduced target gene set. Reason for reducing the number of target genes of the AffymetrixU133P2.0 pathway models was the intended use of pathway analysis on formalin fixed paraffin embedded tissue (FFPE). To avoid multiplexing of qPCRs which leads unavoidably to less reliable qPCR performance, multiple separate diagnostic grade qPCRs needed to be developed. Since frequently very small samples are available from biopsy FFPE material, RNA isolated from such samples is limiting with respect to the number of qPCRs that can be performed. For this reason, a smaller selection of target genes exhibiting optimal performance in the Affymetrix models was identified using signaling pathway analysis results of large Affymetrix microarray datasets containing data of a variety of normal and diseased tissue and cell samples (including multiple cancer types from various organ locations) with a known (‘ground truth”) signaling pathway activity. On such Affymetrix microarray datasets the contribution of target genes to a pathway activity score can be quantified, which is an essential part of the selection process. Subsequently this gene selection was validated by comparing data analysis results obtained with the originally described gene set of the Affymetrix model with an Affymetrix model that was adapted to this limited target gene set. Following this successful initial validation step, qPCR assays were developed according to standard procedures for each of the selected target genes and a set of reference genes. The Bayesian computational qPCR models were constructed and calibrated

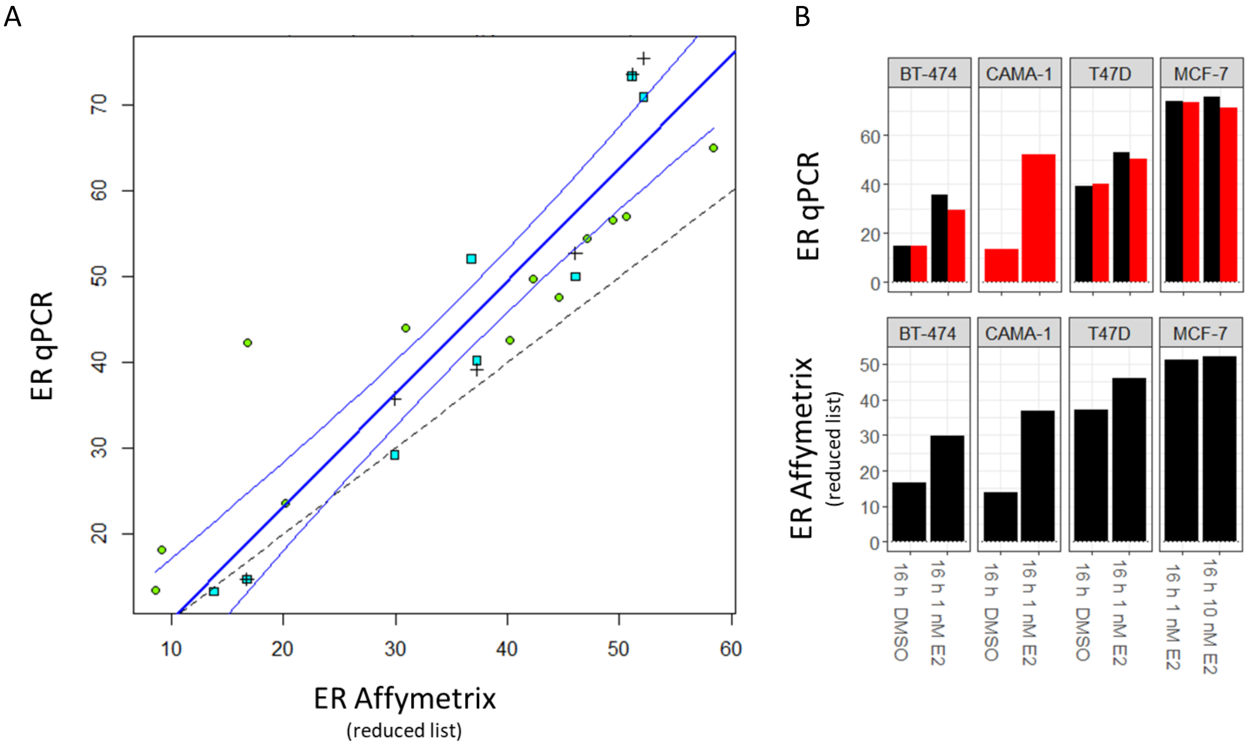


*Figure S-M1. Conversion of the Affymetrix ER pathway model to qPCR pathway model. A. Correlation between results obtained with the Affymetrix U133 P2.0 (reduced target gene list) model and the qPCR model for 12 breast cancer tissue samples (cyan squares, this study) and for 8 breast cancer cell line samples from the GSE127760 public dataset[2] (green squares), 6 samples were re-measured in the qPCR platform (plusses). Dashed line: identity line, blue lines: orthogonal fitting and 95% confidence interval. Affymetrix and qPCR activity scores are strongly correlated, correlation=0.92, 95% CI=0.83-0.96, p-value<0.001, n=26). B. ER pathway activity scores (Y-axis, log2 odds of pathway activity) for the cell line samples GSE127760. Top: qPCR model, bottom: Affymetrix U133 P2.0 (reduced target gene list). Black: first measurement, red: second measurement.*

using cell line calibration sets as previously described[1–3]. Target gene mRNA expression values from cell line experiments were obtained and used as input to calibrate the PCR models, as described previously for the Affymetrix-model. Validation results for the ER pathway model on breast cancer cell line and tissue samples are shown as an example of this procedure (Figure S-M1). Signaling pathway activity scores are reported on a 0-100 scale, computed by normalizing the log2 odds of the calculated probability that the corresponding transcription factor was in the active transcribing state (the readout of the Bayesian model is the probability p that the transcription factor is active, which can be transformed to log2 odds via: $\log_2 (p/(1-p))$).

**Sample preparation and pathway analysis: technical noise analysis**

To exclude confounding technical factors in the sample collection, processing, and qPCR measurements, which could potentially have influenced the comparison between variation in signal transduction pathway activity on macro- (block samples) versus micro-scale (quadrant samples) in the primary tumor, an extensive technical noise analysis was performed.

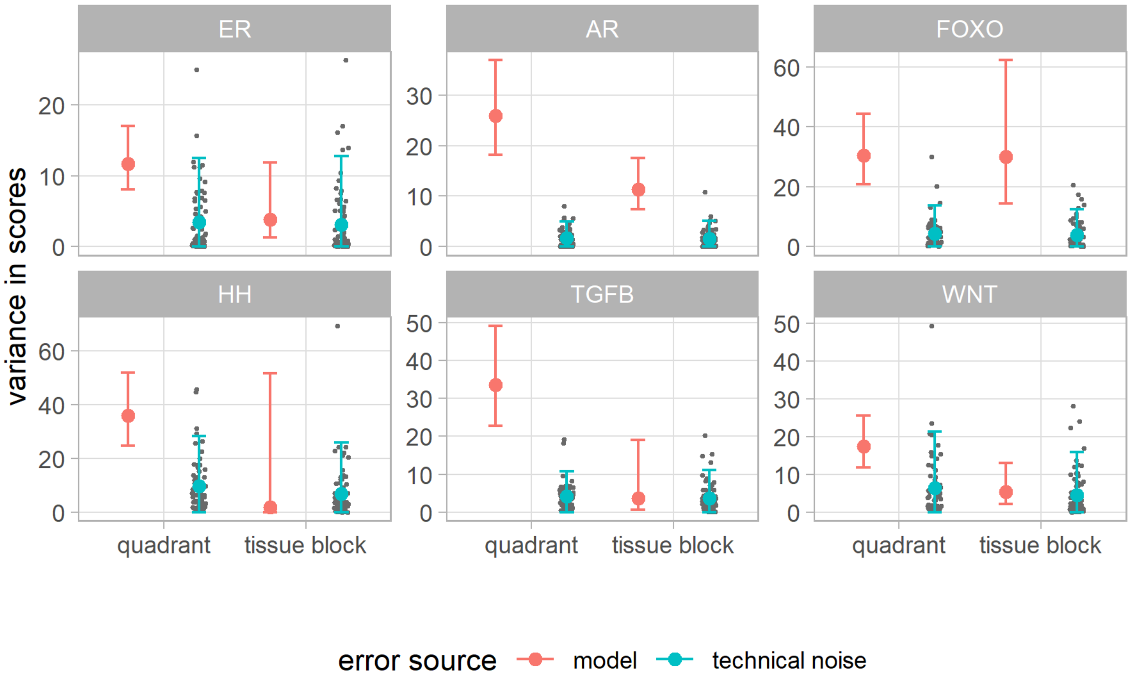


*Figure S-M2. Comparison between the estimated qPCR-related technical noise variance (blue) in relation to the model variance (red) as estimated by the statistical model, including 95% confidence interval (see Figure 3B). As can be seen, the technical noise is smaller than the variance of signaling pathway activity calculated by the model, indicating true intra-tumor heterogeneity as the main cause of the variance. Vertical scale = variance in pathway activity score. This is the natural unit to consider for this question, as variances of spread sources add up.* ***Grey*** *is the simulated technical noise, providing a different value for each individual measurement (patient – tumor – repeated measurement within tumor).* ***Blue*** *is the technical noise variance mean and 95% CI.* ***Red*** *shows the variances, including 95% confidence interval, estimated by the model (which accounts for the sum of technical noise variation + true intra-tumor heterogeneity, and is assumed to be equal in all cases.*

##### 1. *Technical noise associated with performing qPCR*

Investigated was if the qPCR technical noise was the root cause of the observed variance within the samples. The statistical model assumes the variances from different sources add up; i.e. not only the variance due to tumor heterogeneity (goal of the study) but also the technical noise. Part of the qPCR pathway plate product (FIPA Pathway Plate 1.0 from Molecular Pathway Dx, Philips, Eindhoven, The Netherlands) is a qPCR noise estimation model, which calculates a sample specific confidence interval for the reported pathway score. By comparing the variances computed as qPCR noise to the total statistical model variance, it can be estimated how much of the model variance is due to qPCR technical noise (Figure S-M2). Observed from the statistical model is that for all pathway models the technical noise variance is lower than the model variance. Therefore, we concluded that the variances estimated by the statistical model are not caused by qPCR noise only and additional variances, like tumor heterogeneity, are present in our dataset.

##### 2. *High reference gene Cq values*

To exclude that increased technical noise due to high reference gene Cq values caused a bias by impacting reference gene mRNA measurements, additional analysis was performed by eliminating all sample data with an average reference gene Cq value above either 31 or 32

(Figure S-M3). Removing high Cq samples (for sample numbers, see Supplemental Sample Counts) resulted in a small shift in the ratio between primary tumor macro- and micro-heterogeneity, reducing FOXO activity variation on the microscale, while eliminating significant differences between macro and micro-scale variation for AR and Wnt pathways. However, the higher variation at the micro-level for the Hedgehog and TGFβ pathways remained clearly present, while variation in PI3K-FOXO pathway activity increased at the macroscale. As a consequence of this part of the noise analysis, we concluded that some technical bias may have been introduced because a number of samples had been very small resulting in higher technical noise, and for this reason we focus in the discussion and conclusions on the observations that were consistent, and independent of technical noise. Of note, the number of samples left in the analysis following the described exclusion of high Cq samples was too low to perform independent statistical analysis.

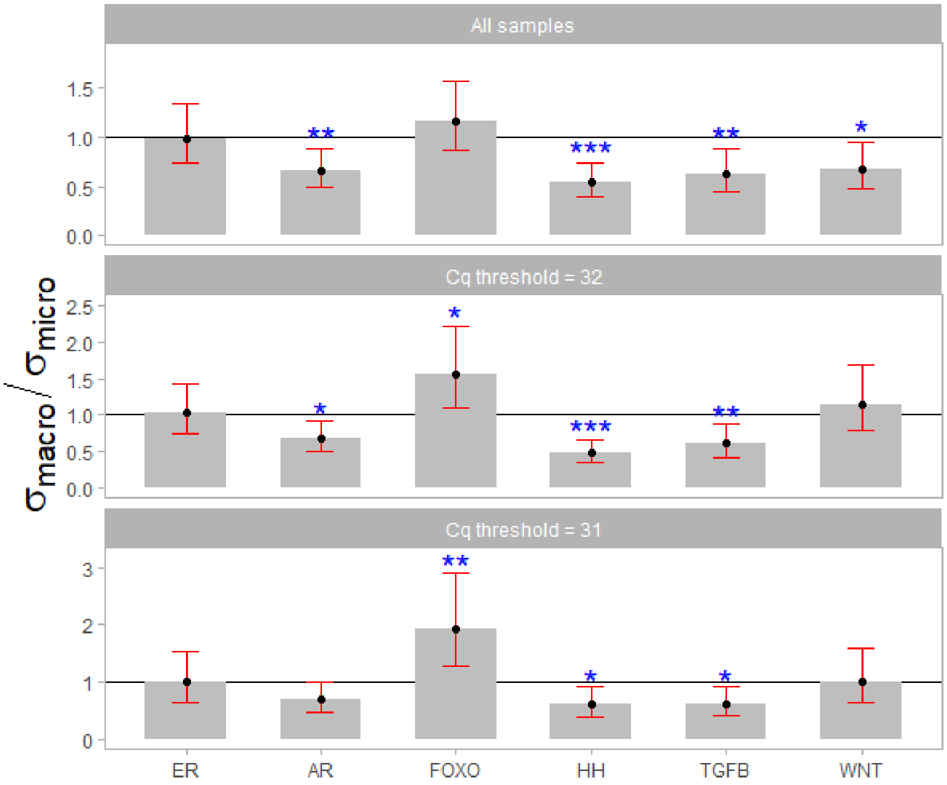


*Figure S-M3. Ratio between macro-scale and micro-scale standard deviation (SD) of signaling pathway activity scores ($\sigma_{macro}/\sigma_{micro}$, measure in tissue block samples and quadrant samples, respectively, similar to Figure 5D). Top: all samples used, middle: left out all samples with average Cq values > 32 for the reference genes used for the qPCR measurements, bottom: left out all samples with average Cq values > 31 for the reference genes used for the qPCR measurements. Variances computed using the macro vs. micro scale model described in detail in supplementary statistics.* $\sigma_{macro}/\sigma_{micro} = \sqrt{(\sigma^2_{sample,block\ sample} + \sigma^2_{block})}/\sigma_{sample,quadrant\ sample}$ *Significance level of Wald test p-value for comparing the ratio* $\sigma_{macro}/\sigma_{micro}$ *to 1 are indicated by stars (*: p-value <0.05, **: p-value <0.01, ***: p-value <0.001.*

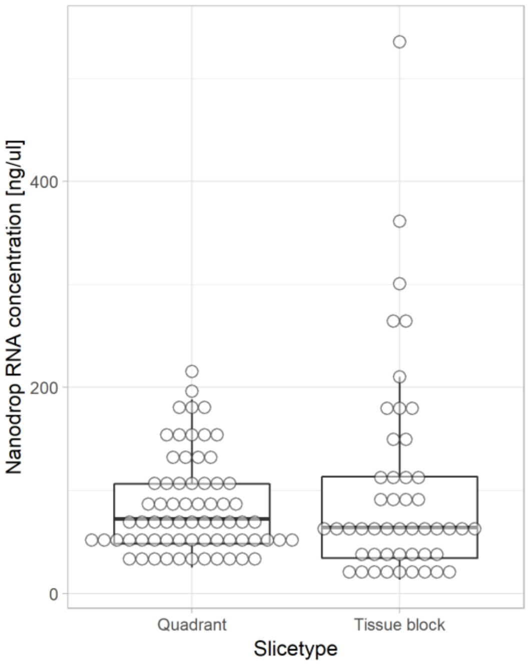


*Figure S-M4. Comparison between amounts of RNA which were isolated from tissue samples, for quadrant and block samples.*

##### 3. *Comparison of tissue sample size between block and quadrant samples*

No significant differences were found between block and quadrant samples with respect to tissue volume from which RNA was isolated, excluding this as a confounding factor (Figure S-M4).

**Deriving PI3K pathway activity from FOXO activity and SOD2 expression**

FOXO transcription factor activity can indicate two functions, either suppression of cell division, or protection against cellular oxidative stress[4]. We have shown before that in Luminal A samples FOXO activity scores are similar to FOXO activity scores in healthy tissue, while low SOD2 expression levels indicate that cellular oxidative stress is absent in Luminal A breast cancer [5]. This is characteristic for the FOXO transcriptional factor being functionally active to control cell division, rather than to protect against cellular oxidative stress [5]. Therefore, Luminal A tissue samples can be used as a surrogate sample set to determine normal FOXO activity scores and SOD2 expression levels. Using the Luminal A breast cancer tissue sample set, the mean of the FOXO activity score minus 2 standard deviations (SD) was defined, as an approximate threshold for FOXO activity, below which FOXO activity levels scores are indicative for activity of the PI3K pathway, and inversely related to activity of this pathway. The established threshold level was 64.1 for the FOXO pathway activity score. Below this threshold, the PI3K pathway was considered as increasingly active and negatively correlated with lower FOXO activity scores. Subsequently, in the FOXO-active Luminal A sample set, the mean SOD2 gene expression level was calculated and the lower threshold for oxidative stress defined as 2 SD above the mean. In this way, -2.03 log2 odds expression level was defined as a threshold above which FOXO activity is increasingly likely to reflect a cellular condition in which the function of FOXO is not any more to control cell division but to protect against cellular oxidative stress [5] (Figure S-M5).

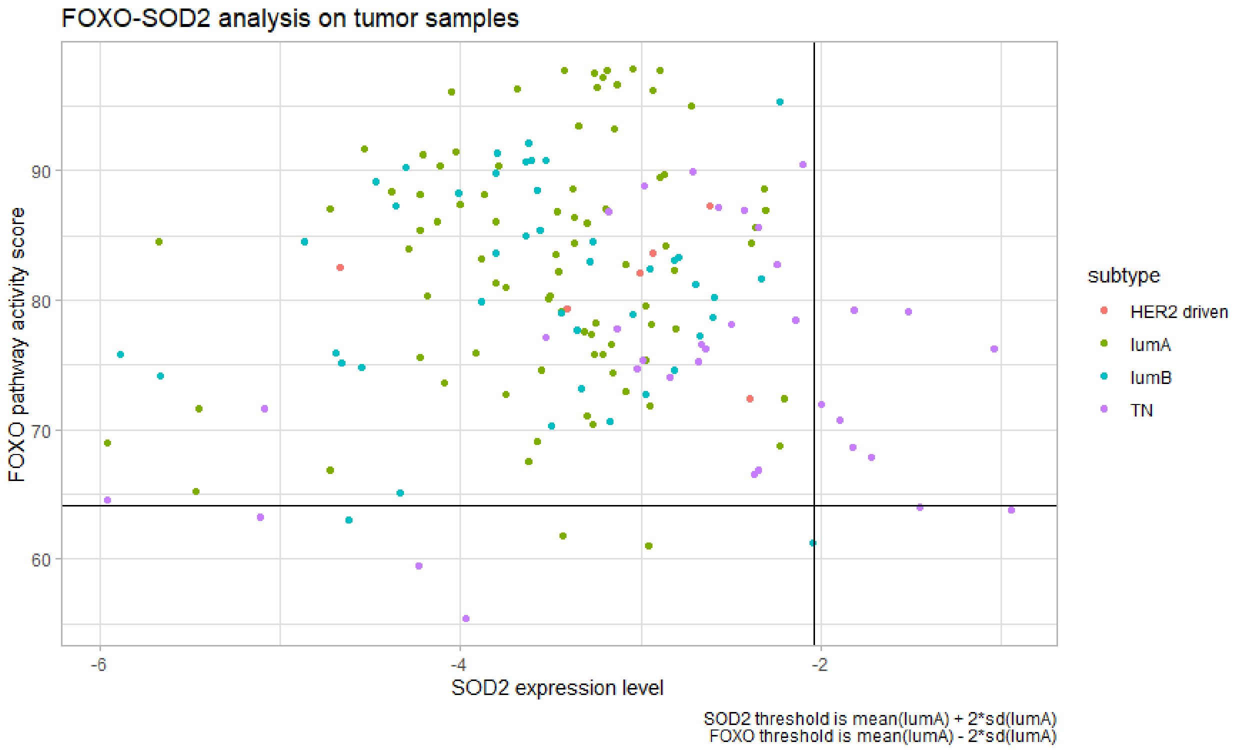


*Figure S-M5. Verification of inverse FOXO-PI3K readout. Samples with high SOD2 expression level could have oxidative stress activate the FOXO pathway, which would mask a positive PI3K readout. Thresholds for risk are determined as shown in the caption.*
