## Supplementary statistics for "Heterogeneity in signaling pathway activity within primary and between primary and metastatic breast cancer"

This document describes the statistical models in more detail, and includes additional statistics not presented in the main text.

Formal model assumptions are that the random effects and residuals follow a normal distribution, at least approximately. In the course of the analysis, the raw data was plotted and no cause for deviating from this assumption was found.

For several of the methods below we used linear mixed models. They can be thought of as a variant of nested ANOVA. Alternatively, they can be thought of as regression where intercepts are random. Linear mixed models are often used in repeated measures ANOVA, e.g. where each patients contributes several values; here the intercept would have a random contribution from each individual patient. We use a variant of the linear mixed models where the standard deviation parameter is allowed to vary between subgroups, whereas usually it is assumed constant. For instance, when comparing the heterogeneity in pathway activity scores between breast cancer subtypes (section 1), we would like to know if the variance in pathway activity scores from block samples from the same patient is dependent the breast cancer subtype, therefore the standard deviation $SD(\beta_{pb,t}) = \sigma_{block,t}$ is subtype dependent. Linear mixed models are often used to study *fixed* effects, where the random effects merely serve to get a reasonable model of the dependencies in the data to ensure that inference on fixed effects is correct. However, the main interest in this study lies in the size of *random* effects, as measured by standard deviations and functions of these.

The mixed linear model algorithms calculate quantities on a $\ln$ scale The *raw* values are therefore log transformed and need to be transformed back (using the exponential function) to obtain the quantities that are being modelled. Computations to obtain p values and confidence intervals are also done in this logarithmic scale. Generally, we use Wald tests to obtain p values and approximate 95% confidence intervals are constructed by subtracting and adding 1.96 standard errors to the point estimated and then transformed back to obtain the desired values. Modeling and statistical analysis were performed in Stata 15.1 and R (nlme, emmeans packages).

### 1. Heterogeneity between subtypes

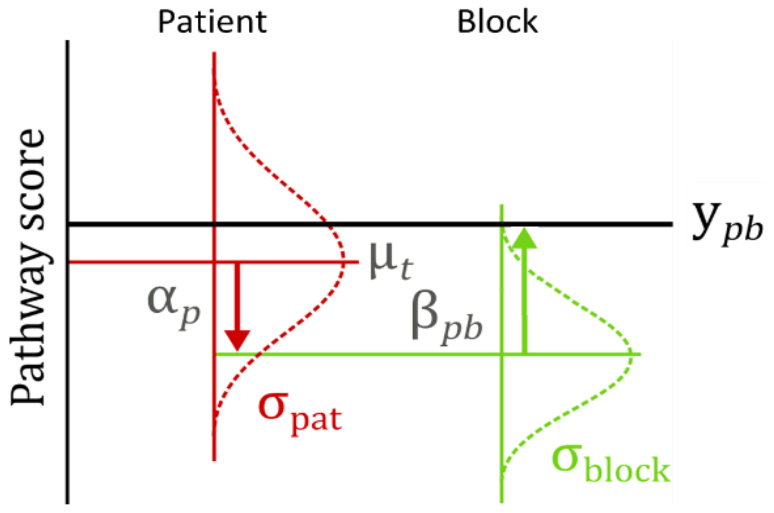


$$y_{pb,t} = \mu_t + \alpha_p + \beta_{pb,t}$$

$$\sigma_{\text{block},t} = SD(\beta_{pb,t})$$

*Figure S-S1: Diagram of heterogeneity between subtype classification models*

Figure S-S1 presents the diagram of the model used to answer questions related to heterogeneity between breast cancer subtypes. This model assumes that the pathway activity score can be built up as a linear combination of variables.

- $y_{pb,t}$: The pathway activity score measured for patient $p$ and (FFPE) tissue block $b$. The sub-index $t$ indicates the patient's breast cancer subtype, with $t$ = Lum A, Lum B or ER– (TN/HER2)
- $\mu_t$: The average activity scores within breast cancer subtype $t$, with $t$ = Lum A, Lum B, or ER–.
- $\alpha_p$: This is the patient specific deviation from the average score $\mu_t$.
    - The standard deviation over all patient contributions $\alpha_p$ is $\sigma_{\text{pat}}$, it indicates how the pathway score can vary between patients and this parameter does not depend on the breast cancer subtype the patient belongs to.
- $\beta_{pb}$: The FFPE block specific contribution attributed to FFPE block $b$ for a particular patient $p$. For each patient tumor, multiple FFPE blocks were available, and this $\beta_{pb}$ parameter is the block specific deviation from the patient's average value $\mu_t + \alpha_p$.
    - Here, three standard deviations were estimated depending on the breast cancer subtype $t$: $\sigma_{\text{block,Lum A}}$ is the spread of pathway scores between block samples for patients with Luminal A breast cancer, $\sigma_{\text{block,Lum B}}$ is the spread of pathway scores between block samples for patients with Luminal B breast cancer, and $\sigma_{\text{block,ER}-}$ is the spread of pathway scores between block samples for patients with ER negative (TN/HER2) breast cancer.

Table S-S1 presents results of pairwise tests for equality of means for $\mu_{\mathrm{ER-}}$, $\mu_{\mathrm{Lum\ A}}$, and $\mu_{\mathrm{Lum\ B}}$ and Tables S-S2 and S-S3 present estimate values for the model parameters. Values were computed using the data from the tissue block samples of sample sets I and II.

The p-values from the log likelihood ratio tests between the subtype dependent $\sigma_{\mathrm{block,t}}$ model and corresponding subtype independent $\sigma_{\mathrm{block}}$ model are all above 0.1 (Table S-S3). This indicates that the variation in activity score between tumors is comparable for all the subtypes and that we may use the simpler subtype independent $\sigma_{\mathrm{block}}$ model in our (further) analysis. The similarity between the two model types can be checked by comparing the average estimates ($\mu_{\mathrm{t}}$) and the standard deviation estimates ($\sigma_{\mathrm{pat}}$) computed using the subtype dependent $\sigma_{\mathrm{block,t}}$ model (Tables S-S2 and S-S3, respectively) and using subtype independent $\sigma_{\mathrm{block}}$ model (Tables S-S4 and S-S5, respectively). Table S-S5 also presents estimates for the ratios $\sigma_{pat}/\sigma_{block}$ and Wald test p-values for comparing the ratios $\sigma_{pat}/\sigma_{block}$ to 1.

*Table S-S1: Pairwise test for equality of means for $\mu_{LumA}$, $\mu_{LumB}$, and $\mu_{ER-}$ computed using the between subtype heterogeneity model, as defined in Figure S-S1. est (lb, ub); point estimate of the difference and 95% confidence interval, p value: Adusted Wald's test p-value. Confidence intervals and p-values adjusted with Tukey method for comparing a family of 3 estimates using R emmeans package.*

| **Pathway** | **test** | **est (lb, ub)** | **p value** | |
|---|---|---|---|---|
| ER | $\mu_{\mathrm{Lum\ A}} - \mu_{\mathrm{Lum\ B}} = 0$ | 1.6 (-6.9, 10.2) | 0.9 | |
| | $\mu_{\mathrm{Lum\ A}} - \mu_{\mathrm{ER-}} = 0$ | 30.4 (22, 38.8) | <0.0001 | *** |
| | $\mu_{\mathrm{Lum\ B}} - \mu_{\mathrm{ER-}} = 0$ | 28.8 (19.1, 38.5) | <0.0001 | *** |
| AR | $\mu_{\mathrm{Lum\ A}} - \mu_{\mathrm{Lum\ B}} = 0$ | 1.4 (-5.2, 8) | 0.9 | |
| | $\mu_{\mathrm{Lum\ A}} - \mu_{\mathrm{ER-}} = 0$ | -0.7 (-7.6, 6.1) | 1 | |
| | $\mu_{\mathrm{Lum\ B}} - \mu_{\mathrm{ER-}} = 0$ | -2.2 (-9.8, 5.5) | 0.8 | |
| FOXO | $\mu_{\mathrm{Lum\ A}} - \mu_{\mathrm{Lum\ B}} = 0$ | 3.0 (-3.9, 9.9) | 0.5 | |
| | $\mu_{\mathrm{Lum\ A}} - \mu_{\mathrm{ER-}} = 0$ | 8.2 (0.8, 15.7) | 0.03 | * |
| | $\mu_{\mathrm{Lum\ B}} - \mu_{\mathrm{ER-}} = 0$ | 5.2 (-2.9, 13.3) | 0.3 | |
| HH | $\mu_{\mathrm{Lum\ A}} - \mu_{\mathrm{Lum\ B}} = 0$ | 2.8 (-0.4, 6.1) | 0.09 | . |
| | $\mu_{\mathrm{Lum\ A}} - \mu_{\mathrm{ER-}} = 0$ | -0.6 (-4.3, 3.2) | 0.9 | |
| | $\mu_{\mathrm{Lum\ B}} - \mu_{\mathrm{ER-}} = 0$ | -3.4 (-7.4, 0.6) | 0.1 | |
| TGFb | $\mu_{\mathrm{Lum\ A}} - \mu_{\mathrm{Lum\ B}} = 0$ | 5.1 (-3, 47) | 0.3 | |
| | $\mu_{\mathrm{Lum\ A}} - \mu_{\mathrm{ER-}} = 0$ | 8.3 (0.3, 16.4) | 0.04 | * |
| | $\mu_{\mathrm{Lum\ B}} - \mu_{\mathrm{ER-}} = 0$ | 3.3 (-6, 12.5) | 0.7 | |
| WNT | $\mu_{\mathrm{Lum\ A}} - \mu_{\mathrm{Lum\ B}} = 0$ | 4.7 (-2.5, 11.9) | 0.3 | |
| | $\mu_{\mathrm{Lum\ A}} - \mu_{\mathrm{ER-}} = 0$ | -5.2 (-12.5, 2.2) | 0.2 | |
| | $\mu_{\mathrm{Lum\ B}} - \mu_{\mathrm{ER-}} = 0$ | -9.9 (-18.1, -1.6) | 0.02 | * |

*Table S-S2: Average activity score ($\mu_t$) estimates for the subtype dependent $\sigma_{block,t}$ model (as defined in Figure S-S1) for each signaling pathway. Average scores presented as point estimate (95% confidence interval).*

| Pathway | $\boldsymbol{\mu}_{\textbf{Lum A}}$ | $\boldsymbol{\mu}_{\textbf{Lum B}}$ | $\boldsymbol{\mu}_{\textbf{ER}-}$ |
|---|---|---|---|
| ER | 55.8 (51.7, 60) | 54.2 (48.4, 59.9) | 25.4 (19.8, 31) |
| AR | 30.6 (27.3, 33.9) | 29.2 (24.8, 33.5) | 31.3 (26.7, 35.9) |
| FOXO | 81.7 (78.1, 85.3) | 78.7 (74.3, 83.2) | 73.5 (68.5, 78.6) |
| HH | 22.3 (20.6, 24) | 19.4 (17.3, 21.5) | 22.8 (20.2, 25.4) |
| TGFb | 63.6 (59.7, 67.4) | 58.5 (53.1, 63.9) | 55.3 (49.8, 60.7) |
| WNT | 13.2 (9.6, 16.8) | 8.5 (3.7, 13.3) | 18.4 (13.5, 23.2) |

*Table S-S3: Standard deviation ($\sigma_{pat}$ and $\sigma_{block,t}$) estimates for the subtype dependent $\sigma_{block,t}$ model (as defined in Figure S-S1) for each signaling pathway. Standard deviations presented as point estimate (95% confidence interval), p-value: p-value from log likelihood ratio tests between subtype dependent $\sigma_{block,t}$ model and corresponding subtype independent $\sigma_{block}$ model computed with anova.lme function from nlme R package.*

| Pathway | $\boldsymbol{\sigma}_{\textbf{pat}}$ | $\boldsymbol{\sigma}_{\textbf{block,Lum A}}$ | $\boldsymbol{\sigma}_{\textbf{block,Lum B}}$ | $\boldsymbol{\sigma}_{\textbf{block,ER}-}$ | p-value |
|---|---|---|---|---|---|
| ER | 8.0 (6.1, 10.5) | 3.9 (3.1, 5.0) | 4.7 (3.3, 6.6) | 2.9 (2.0, 4.1) | 0.2 |
| AR | 6.1 (4.6, 8.2) | 4.0 (3.1, 5.2) | 3.2 (2.3, 4.4) | 4.3 (3.0, 6.3) | 0.4 |
| FOXO | 5.6 (3.8, 8.2) | 7.4 (5.8, 9.5) | 5.8 (4.2, 8.0) | 7.7 (5.5, 10.8) | 0.4 |
| HH | 2.5 (1.6, 3.9) | 3.7 (2.9, 4.7) | 2.8 (2.0, 3.9) | 4.5 (3.3, 6.2) | 0.1 |
| TGFb | 7.1 (5.3, 9.6) | 4.8 (3.8, 6.1) | 6.1 (4.5, 8.4) | 5.8 (4.0, 8.4) | 0.5 |
| WNT | 6.9 (5.3, 9.0) | 3.3 (2.6, 4.2) | 2.6 (1.9, 3.6) | 3.0 (2.1, 4.3) | 0.5 |

*Table S-S4: Average activity score ($\mu_t$) estimates for subtype dependent $\sigma_{block}$ model for each signaling pathway. Average scores presented as point estimate (95% confidence interval).*

| Pathway | $\boldsymbol{\mu}_{\textbf{Lum A}}$ | $\boldsymbol{\mu}_{\textbf{Lum B}}$ | $\boldsymbol{\mu}_{\textbf{ER}-}$ |
|---|---|---|---|
| ER | 55.8 (51.6, 60) | 54.2 (48.4, 59.9) | 25.4 (19.7, 31.2) |
| AR | 30.6 (27.2, 33.9) | 29.2 (24.6, 33.7) | 31.3 (26.8, 35.9) |
| FOXO | 81.7 (78.2, 85.3) | 78.7 (74, 83.5) | 73.5 (68.6, 78.4) |
| HH | 22.3 (20.6, 23.9) | 19.4 (17.1, 21.6) | 22.8 (20.5, 25.1) |
| TGFb | 63.6 (59.6, 67.5) | 58.5 (53.2, 63.9) | 55.2 (49.8, 60.7) |
| WNT | 13.2 (9.6, 16.8) | 8.5 (3.6, 13.4) | 18.4 (13.5, 23.3) |

*Table S-S5: Standard deviation ($\sigma_{pat}$ and $\sigma_{block}$) estimates and $\sigma_{pat}/\sigma_{block}$ ratio estimates for the subtype independent $\sigma_{block}$ model, for each signaling pathway. Standard deviations and ratios presented as point estimate (95% confidence interval), p-value: Wald test p-value for comparing the ratio $\sigma_{pat}/\sigma_{block}$ to 1.*

| Pathway | $\boldsymbol{\sigma}_{\textbf{pat}}$ | $\boldsymbol{\sigma}_{\textbf{block}}$ | $\boldsymbol{\sigma}_{\textbf{block}}/\boldsymbol{\sigma}_{\textbf{block}}$ | p-value |
|---|---|---|---|---|
| ER | 8.1 (6.2, 10.5) | 3.9 (3.3, 4.7) | 2.1 (1.5, 2.8) | 0.03 |
| AR | 6.2 (4.7, 8.3) | 3.9 (3.3, 4.6) | 1.6 (1.1, 2.3) | 0.2 |
| FOXO | 5.6 (3.8, 8.3) | 7.1 (6.0, 8.4) | 0.8 (0.5, 1.2) | 0.6 |
| HH | 2.4 (1.5, 3.9) | 3.7 (3.2, 4.4) | 0.6 (0.4, 1.1) | 0.4 |
| TGFb | 7.1 (5.3, 9.6) | 5.4 (4.6, 6.4) | 1.3 (0.9, 1.9) | 0.5 |
| WNT | 6.9 (5.3, 9.0) | 3.0 (2.5, 3.6) | 2.3 (1.7, 3.2) | 0.01 |

To better understand how much each model parameter affects the activity score of each signaling pathway we also carried out a variance components analysis. Such analysis enables us to estimate the amount (or percentage) of variance that can be attributed to (explained by) each (random contribution) term of the model. Higher explained percentages indicate a higher contribution to the total spread in the activity score. Since we are also in interested in the amount of variance explained by the breast cancer subtype the tumor belongs to, we replaced the subtype dependent average estimates ($\mu_t$) by a subtype independent average estimate ($\mu$) plus a subtype specific random contribution ($\gamma_t$) as depicted in Figure S-S2.

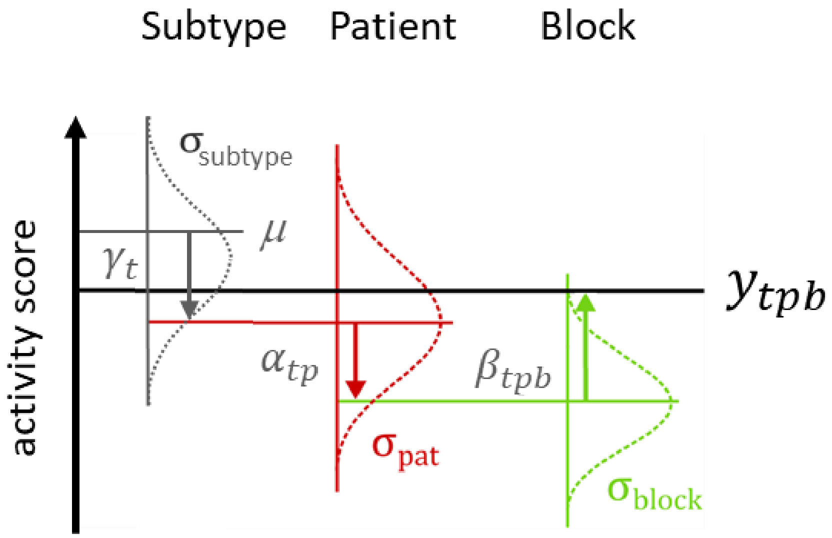


$$y_{tpb} = \mu + \gamma_t + \alpha_{tp} + \beta_{tpb}$$

*Figure S-S2: Modified diagram of heterogeneity between subtype classification models used for variance components analysis. De term $\mu_t$ is replaced by $\mu + \gamma_t$.*

#### 2. Heterogenaity between macro- and micro-scale

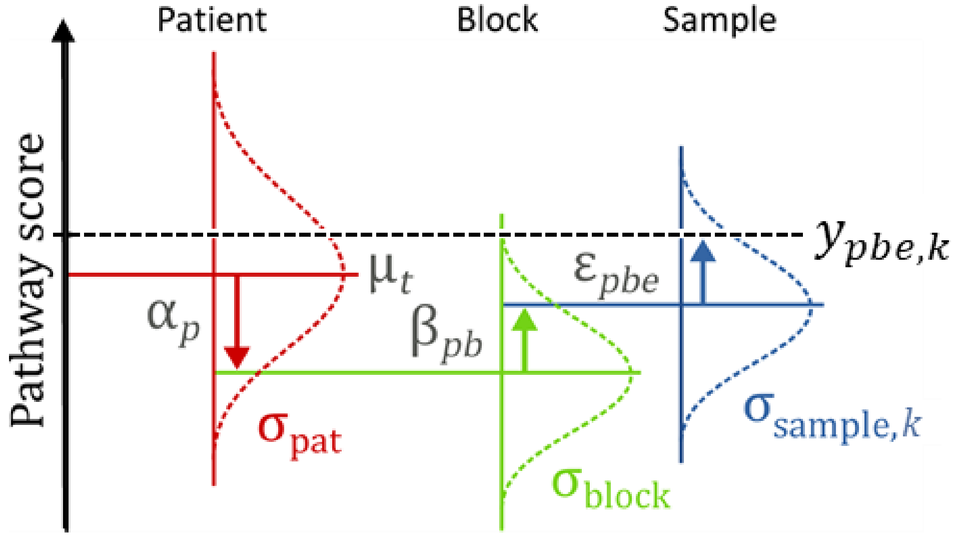


$$y_{pbe,k} = \mu_t + \alpha_p + \beta_{pb} + \varepsilon_{pbe,k}$$

(Tissue block samples heterogeneity)

$$\sigma_{\text{macro}} = \sqrt{\sigma^2_{\text{block}} + \sigma^2_{\text{sample,tissue block sample}}}$$
$$= \sqrt{SD(\beta_{pb})^2 + SD(\epsilon_{pbe,\text{block sample}})^2}$$

(Quadrant samples heterogeneity)

$$\sigma_{\text{micro}} = \sigma_{\text{sample,quadrant sample}}$$
$$= SD(\epsilon_{pbe,\text{quadrant sample}})$$

*Figure S-S3: Diagram of the macro and micro scale model implementing linear mixed models with three layers.*

The macro vs. micro scale modelling approach (Figure S-S3) is very similar to the model explained in section 1 (above). In this model, an additional contribution $\epsilon_{pbe}$ (the blue arrow) is added to the pathway activity score of each block, to model the possibility of quadrant-to-quadrant heterogeneity. Furthermore, the standard deviation of the FFPE block specific contribution ($\sigma_{\text{block}}$) is now independent of breast cancer subtype.

- $y_{pbe,k}$ : The pathway score measured for patient $p$, FFPE block $b$ and sample $e$. The sub-index $k$ indicates the sample type used with $k$ = quadrant sample or tissue block sample, cf. Figure 1 main text.
- $\mu_t$ : The average scores within breast cancer subtype, with $t$ = Lum A, Lum B, or ER– (TN/HER2).
- $\alpha_p$ : The patient specific contribution attributed to patient $p$. This is the patient specific deviation from the average score $\mu_t$.

- The standard deviation over all patient contributions $\alpha_p$ is $\sigma_{\text{pat}}$, it indicates how the pathway score can vary between patients and this parameter does not depend on breast cancer subtype the patient belongs to.

- $\beta_{pb}$ : The FFPE block specific contribution attributed to FFPE block $b$ for a particular patient $p$. For each patient, multiple FFPE blocks were available, and this $\beta_{pb}$ parameter is the block specific deviation from the patient contribution $\alpha_p$.
  - The standard deviation over $\beta$ is $\sigma_{\text{block}}$, and it describes the spread of pathway scores between the FFPE tissue block samples.
- $\varepsilon_{pbe,k}$ : The sample specific contribution attributed to a sample $e$ of type $k$, with $k$ = quadrant sample or (tissue) block sample, for a particular FFPE block $b$ of patient $p$. This $\varepsilon_{pbe,k}$ parameter is the sample specific deviation from the FFPE block contribution $\beta_{pb}$.
  - Here, two different standard deviations $\sigma_{\text{sample},k}$ were computed depending on the sample type $k$: $\sigma_{\text{sample,quadrant sample}}$ is the spread of pathway scores between quadrant samples and $\sigma_{\text{sample,block sample}}$ is the spread of pathway scores between the tissue block samples. This approach of using of two different standard deviations depending on the sample type allows the model to estimate macro-scale and micro-scale tumor heterogeneity, despite the unbalanced design, as described below.

The standard deviation between quadrant samples, $\sigma_{\text{sample,quadrant sample}}$, is a straightforward measure of micro-scale heterogeneity, as term $\epsilon_{pbe,\text{quadrant sample}}$ describes the differences between the four quadrant measurements within a FFPE block (i.e. samples that are spatially close together), therefore $\sigma_{micro} = \sigma_{\text{sample,quadrant samples}}$. The measure for macro-scale heterogeneity is a slightly more complex derive, since it should measure the variation in pathway activity scores between the FFPE blocks within a specific tumor. This encompasses the sum of variances attributed to the FFPE block specific contribution $\beta_{pb}$ and the tissue block sample specific contribution$\epsilon_{pbe,\text{block sample}}$, i.e.:

$$\sigma^2_{\text{macro}} = \sigma^2_{\text{block}} + \sigma^2_{\text{sample,block sample}}, \text{ therefore}$$
$$\sigma_{\text{macro}} = \sqrt{\sigma^2_{\text{block}} + \sigma^2_{\text{sample,block sample}}}.$$

Tables S-S6 and S-S7 present the values of the model's standard deviation parameters and derived estimates thereof.

*Table S-S6: Macro and micro scale model standard deviations ($\sigma$ parameter) estimates, as defined in Figure S-S3, for each signaling pathway presented point estimate (95% confidence interval).*

| **Pathway** | $\boldsymbol{\sigma_{pat}}$ | | $\boldsymbol{\sigma_{block}}$ | | $\boldsymbol{\sigma_{sample,\text{quadrant sample}}}$ | | $\boldsymbol{\sigma_{sample,\text{block sample}}}$ | |
|---|---|---|---|---|---|---|---|---|
| ER | 8.7 | (5.9, 12.8) | 2.8 | (1.8, 4.1) | 3.4 | (2.8, 4.1) | 2.0 | (1.1, 3.4) |
| AR | 5.2 | (3.5, 7.8) | 0.0 | * | 5.1 | (4.3, 6.1) | 3.4 | (2.71, 4.2) |
| FOXO | 6.7 | (4.3, 10.5) | 3.4 | (1.5, 7.7) | 5.5 | (4.6, 6.7) | 5.5 | (3.8, 7.9) |
| HH | 3.0 | (1.8, 5.2) | 3.0 | (1.9, 4.6) | 6.0 | (5.0, 7.2) | 1.4 | (0.3, 7.2) |
| TGFb | 7.9 | (5.3, 11.8) | 3.0 | (2.1, 4.5) | 5.8 | (4.8, 7.0) | 1.9 | (0.9, 4.4) |
| WNT | 6.8 | (4.6, 10.1) | 1.6 | (0.7, 3.7) | 4.2 | (3.4, 5.1) | 2.3 | (1.5, 3.6) |

* unreliable confidence interval.

*Table S-S7: Macro and micro scale model derived parameters for each signaling pathway presented point estimate (95% confidence interval).* $\sigma_{micro}$: $\sigma_{sample,quadrant\ sample}$, $\sigma_{macro} = \sqrt{(\sigma^2_{block} + \sigma^2_{sample,block\ sample})}$, *p value: Wald test p-value for comparing the ratio* $\sigma_{macro}/\sigma_{micro}$ *to 1.*

| **pathway** | $\boldsymbol{\sigma_{macro}}$ | | $\boldsymbol{\sigma_{micro}}$ | | $\boldsymbol{\sigma_{macro}/\sigma_{micro}}$ | | **p value** |
|---|---|---|---|---|---|---|---|
| ER | 3.4 | (2.7, 4.3) | 3.4 | (2.8, 4.1) | 1.0 | (0.7, 1.3) | 0.9 |
| AR | 3.4 | (2.7, 4.2) | 5.1 | (4.3, 6.1) | 0.7 | (0.5, 0.9) | 0.004 |
| FOXO | 6.4 | (5.2, 8.0) | 5.5 | (4.6, 6.7) | 1.2 | (0.9, 1.6) | 0.3 |
| HH | 3.3 | (2.6, 4.2) | 6.0 | (5.0, 7.2) | 0.5 | (0.4, 0.7) | 0.0001 |
| TGFb | 3.6 | (2.8, 4.6) | 5.8 | (4.8, 7.0) | 0.6 | (0.4, 0.9) | 0.006 |
| WNT | 2.8 | (2.2, 3.6) | 4.2 | (3.4, 5.1) | 0.7 | (0.5, 0.9) | 0.02 |

### 3. Heterogeneity between metastases and matched primary tumors

Figure S-S4 describes the model used for comparing pathway activity scores between primary tumor and matched lymph node (LN) metastasis samples. For this analysis, 8 patients were used for which at least one LN metastasis and multiple primary tumor (PT) tissue block samples were available (matched patients of sample set IV and sample set II[1]; cf. Supplementary Information, SampleCounts and Data). Due to this very low sample count and unbalanced number of samples per patient (for five patients there were one or two lymph node samples, and for the remaining three patients there were between seven and ten samples), many statistical tests are either inconclusive or not very meaningful. For this reason, we used a simpler approach where the scores of the individual metastatic samples are described as the average score of the respective primary tumor (tissue) block samples plus a LN metastatic sample specific contribution:

$$y_{p,\mathrm{LN}_i} = \bar{y}_{p,\mathrm{block}} + \Delta_{p,\mathrm{LN}_i},$$

for each LN metastatic sample $i$ belonging to each patient $p$ with multiple matched PT tissue block samples. Here, $\bar{y}_{p,\mathrm{block}}$ is the mean activity score of the available PT tissue block samples (sample set II) and is computed for each patient separately (Figure S-S4 Left), and the LN metastatic sample specific

[1] Samples of patient 20 not used in this analysis because of only one PT sample was available.

contribution $\Delta_{p,\mathrm{LN}_i}$ is the difference between the activity score of the LN sample and the score of the average score of the respective primary tumor samples.

$$\Delta_{p,\mathrm{LN}_i} = y_{p,\mathrm{LN}_i} - \bar{y}_{p,\mathrm{block}}$$

Similarly, the differences between primary tumor (tissue) block scores and the respective block averages

$$\Delta_{p,\mathrm{block}_j} = y_{p,\mathrm{block}_j} - \bar{y}_{p,\mathrm{block}},$$

(that can be considered as tissue block sample specific contributions), were also computed for each tissue block sample $i$ belonging to each patient $p$ (Figure S-S4 Left).

To investigate if the variation in score between primary tumor and LN metastasis is larger than the variation in score between multiple samples of the same primary tumor, we compared the histograms of the LN metastatic sample specific contributions, $\Delta_{p,\mathrm{LN}_i}$, with histograms of the block sample specific contributions, $\Delta_{p,\mathrm{block}_j}$, as shown for the FOXO model (Figure S-S4 Center).

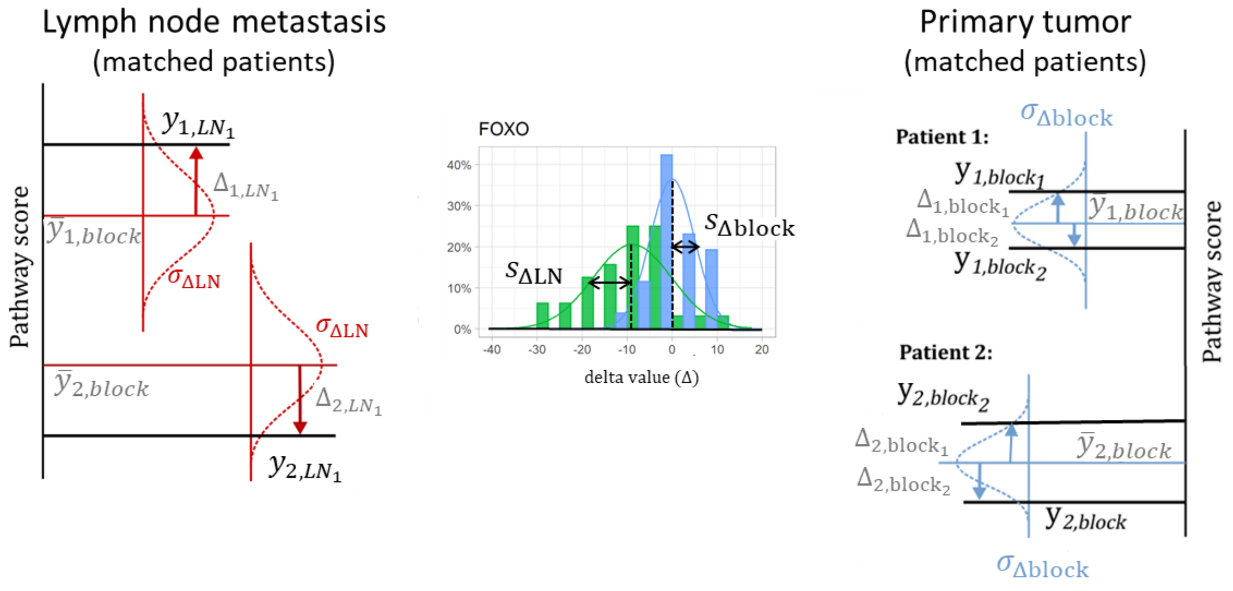


$y_{p,\mathrm{LN}_i} = \bar{y}_{p,block} + \Delta_{p,\mathrm{LN}_i}$ $\qquad$ $y_{p,\mathrm{block}_j} = \bar{y}_{p,block} + \Delta_{p,\mathrm{block}_j}$

*Figure S-S4: Lymph node metastases and matched primary tumor model. Left: diagram of LN metastases model. Right: diagram of primary tumor block model. Center: frequency counts of $\Delta_{LN}$ (green) and $\Delta_{block}$ (blue) approximate distributions of the LN metastatic sample specific contributions and the tissue block specific contributions respectively.*

Figure S-S5 describes the model used for comparing pathway activity scores between primary tumor and matched distant metastasis samples. The model used for this analysis is identical to the one used in the LN metastasis analysis, but in this case only one primary tumor and at least one matched distant metastasis samples were available (Sample set III; cf. Supplementary Information, SampleCounts and Data). Because only one matched primary tumor sample was available per patient, in this case the scores of the individual metastatic samples were described as the score of the respective primary tumor sample plus a metastatic sample specific contribution:

$$y_{q,\mathrm{meta}_k} = y_{q,\mathrm{prim}} + \Delta_{q,\mathrm{meta}_k},$$

for each metastatic sample $k$ belonging to each patient $q$ of sample set III. Here, the activity score of the available primary sample is assumed representative of the respective primary tumor and is computed for each patient separately (Figure S-S5 Left), and the metastatic sample specific contribution $\Delta_{q,\mathrm{meta}_k}$ is the difference between the activity score of the metastatic sample and the score of the respective primary tumor sample.

Here again, because of lack of multiple matched PT sample, the same differences between primary tumor (tissue) block scores and the respective block averages as used for the LN model ($\Delta_{p,\mathrm{block}_j}$) were used as reference (Figure S-S4 Left). And we compared the histograms of the metastatic sample specific contributions, $\Delta_{q,\mathrm{meta}_k}$, with histograms of the block sample specific contributions, $\Delta_{p,\mathrm{block}_j}$, to investigate if the variation is score between primary tumor and metastatic sample was larger than the variation in score between multiple samples of the same primary tumor, (Figure S-S5 Center).

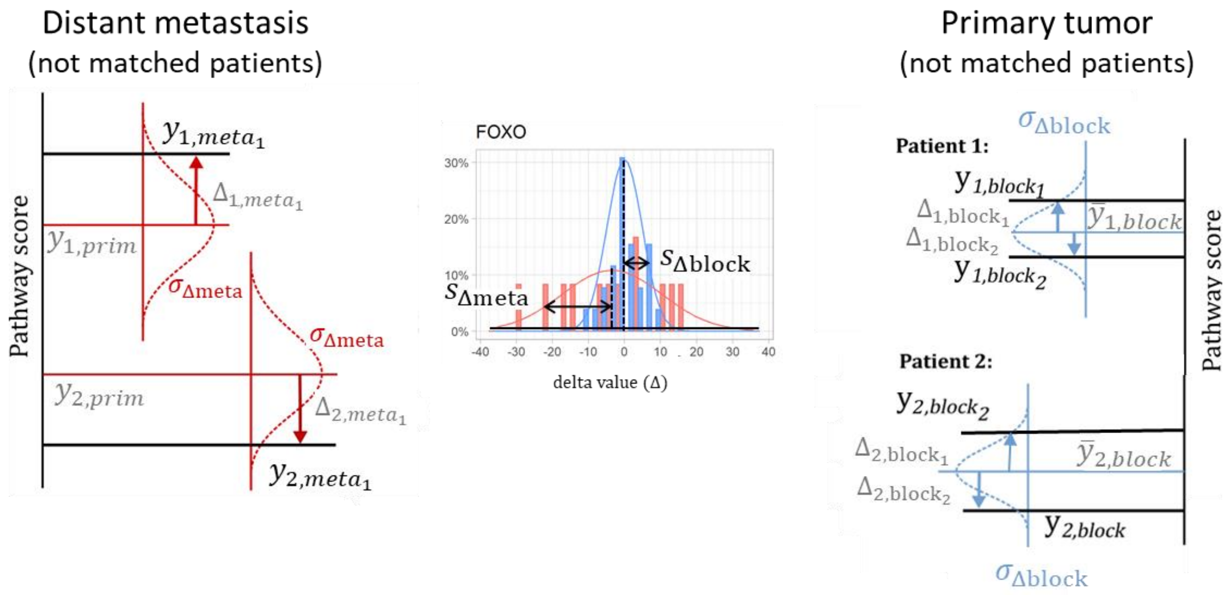


$y_{q,\mathrm{meta}_k} = y_{q,\mathrm{prim}} + \Delta_{q,\mathrm{meta}_k}$ $\quad$ $y_{p,\mathrm{block}_i} = \bar{y}_{p,block} + \Delta_{p,\mathrm{block}_i}$

*Figure S-S5: Distant metastases and matched primary tumor model. Left: diagram of distant metastases and matched primary tumor model. Right: diagram of primary tumor block scores and the respective block averages. Center: frequency counts of $\Delta_{meta}$ (red) and $\Delta_{block}$ (blue) approximate distributions of the metastatic sample specific contribution and the tissue block specific contributions respectively.*

The primary sample activity scores $y_{p,\mathrm{prim}}$ (metastatic sample set III. Figure S-S5 Left) were computed based on a single sample, while the mean activity scores $\bar{y}_{q,\mathrm{block}}$ (primary tumor sample set II, Figure S-S5 Right) were computed based on the average of multiple samples. Because of these different approaches, we expect that the metastatic sample specific contributions, $\Delta_{q,\mathrm{meta}_k}$, will be typically larger than the block sample specific contributions, $\Delta_{p,\mathrm{block}_j}$, due to the different way the error propagates in the two cases:

For the metastatic patient samples

$$\Delta_{q,\mathrm{meta}_k} = y_{q,\mathrm{meta}_k} - y_{q,\mathrm{prim}}$$

and the respective standard deviation is

$$s_{\Delta q,meta_k} = \sqrt{s^2_{y_{q.meta_k}} + s^2_{y_{q,prim}}}.$$

Assuming that the technical measurement error noise is the same for all measurements, we have $s^2_{y_{q.meta_k}} = s^2_{y_{q.prim}} = s^2_{\mathrm{y}}$ we have

$$s_{\Delta meta} = s_{\Delta q,meta_k} = \sqrt{s_y^2 + s_y^2} = s_y \cdot \sqrt{2}.$$

For the primary tumor tissue block samples

$$\Delta_{p,\mathrm{block}_j} = y_{p,\mathrm{block}_j} - \bar{y}_{p,\mathrm{block}}.$$

The assumption that the technical measurement error noise is the same for all measurements gives

$$s^2_{y_{p.block_j}} = s^2_{\mathrm{y}},\ \text{and}\ \ s^2_{\bar{y}_{p,block}} = \frac{1}{N^2} s_y^2$$

and the respective primary tumor tissue block sample standard deviation becomes

$$\mathrm{s}_{\Delta\mathrm{block}} = \mathrm{s}_{\Delta p,\mathrm{block}_j} = \sqrt{s_{\mathrm{y}}^2 + \frac{1}{N^2} s_y^2} = s_y \cdot \sqrt{1 + \frac{1}{N^2}}$$

Where $N$ is the number of available tissue block samples for patient $p$.

Since the number of primary tumor tissue block samples differs per patient, we use its average, i.e. $N \approx 2.94$, to estimate the expected ratio

$$\frac{s_{\Delta meta}}{s_{\Delta block}} = \frac{\sqrt{2}\, s_y}{\sqrt{1 + \frac{1}{N}}\, s_y} \approx 1.22$$

From this, an increase in spread of about 22% is expected solely because there is only a single primary tumor sample available for the metastases. A larger spread than this 22% increase could be due to other causes, i.e. the metastatic tumor being different from the primary tumor.

The results of both analysis are presented below. Table S-S8 summarizes the mean and standard deviations obtained for $\Delta_{p,\mathrm{block}_j}$, $\Delta_{p,\mathrm{LN}_i}$and $\Delta_{q,\mathrm{meta}_k}$. Table S-S9 presents corresponding differences between means and ratios between standard deviations for the various pathway models. Figure S-S6 shows the percent frequency counts of the Lymph node metastasis sample specific contributions, $\Delta_{p,\mathrm{LN}_i}$, and of the primary block tissue sample specific contributions, $\Delta_{p,\mathrm{block}_j}$, for the various models. Figure S-S7 show the percent frequency counts of the metastatic sample specific contributions, $\Delta_{q,\mathrm{meta}_k}$, and of the primary block tissue sample specific contributions, $\Delta_{p,\mathrm{block}_j}$, for the various models.

*Table S-S8: Mean and standard deviation computed for $\Delta_{q,block_j}$, $\Delta_{q,LN_i}$, and $\Delta_{p,meta_k}$. Values are given as point estimate and 95%CI.*

| | Primary tumor $\Delta_{q,\mathrm{block}}$ | | Lymph node metastases $\Delta_{q,\mathrm{LN}}$ | | Distant metastases $\Delta_{p,\mathrm{meta}}$ | |
|---|---|---|---|---|---|---|
| | av | (sd) | av | (sd) | av | (sd) |
| ER | 0.00 | (3.01) | -1.89 | (9.38) | -5.88 | (15.2) |
| AR | 0.00 | (3.84) | 4.43 | (7.29) | -3.08 | (11.86) |
| FOXO | 0.00 | (5.12) | -8.92 | (9.01) | -3.47 | (14.45) |
| HH | 0.00 | (2.19) | 1.99 | (5.66) | 4.39 | (9.32) |
| TGFB | 0.00 | (3.44) | -7.02 | (8.26) | -5.93 | (9.03) |
| WNT | 0.00 | (2.23) | 0.33 | (6.49) | 6.77 | (13.06) |

*Table S-S9: differences between means and ratios between standard deviations for the various pathway models.*

| pathway | av(LN) - av(block) | $\frac{s_{\Delta LN}}{s_{\Delta block}}$ | av(meta) - av(block) | $\frac{s_{\Delta meta}}{s_{\Delta block}}$ | av(meta) - av(LN) | $\frac{s_{\Delta meta}}{s_{\Delta LN}}$ |
|---|---|---|---|---|---|---|
| ER | -1.89 | 3.12 | -5.88 | 5.05 | -3.99 | 1.62 |
| AR | 4.43 | 1.90 | -3.08 | 3.09 | -7.50 | 1.63 |
| FOXO | -8.92 | 1.76 | -3.47 | 2.82 | 5.46 | 1.60 |
| HH | 1.99 | 2.58 | 4.39 | 4.25 | 2.40 | 1.65 |
| TGFB | -7.02 | 2.40 | -5.93 | 2.62 | 1.09 | 1.09 |
| WNT | 0.33 | 2.91 | 6.77 | 5.85 | 6.44 | 2.01 |

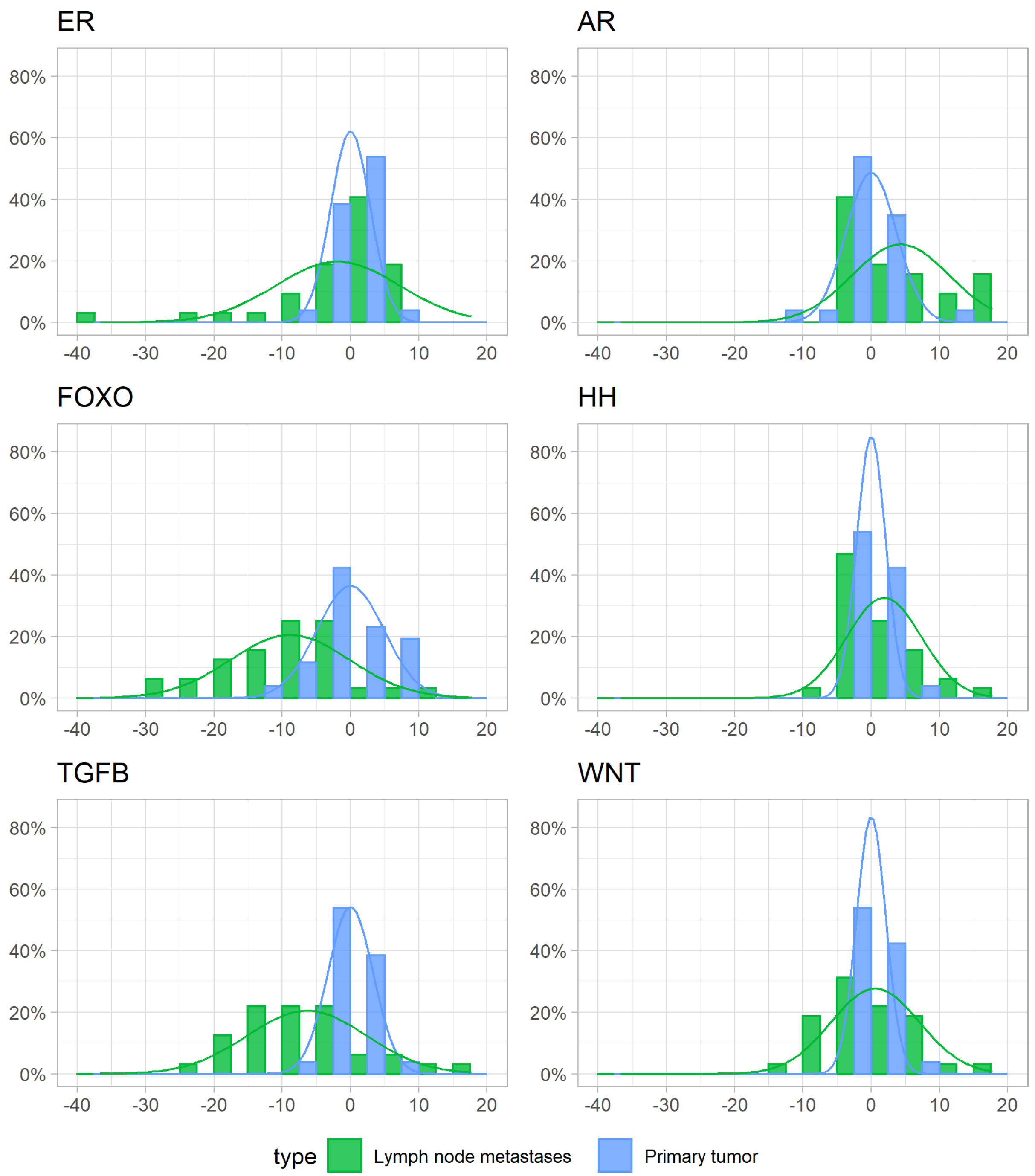


*Figure S-S6: Comparison of frequency counts of the lymph node metastasis sample specific contributions, $\Delta_{p,LN_i}$ (in green), and of the corresponding primary block tissue sample specific contributions, $\Delta_{p,block_j}$ (in blue), for the various signaling pathway models. The bars show the percent frequency counts. As a visual guide, solid lines are added representing Gaussian curves with mean and standard deviation equal to the mean and standard deviation values computed for , $\Delta_{p,LN_i}$ and $\Delta_{p,block_j}$, respectively.*

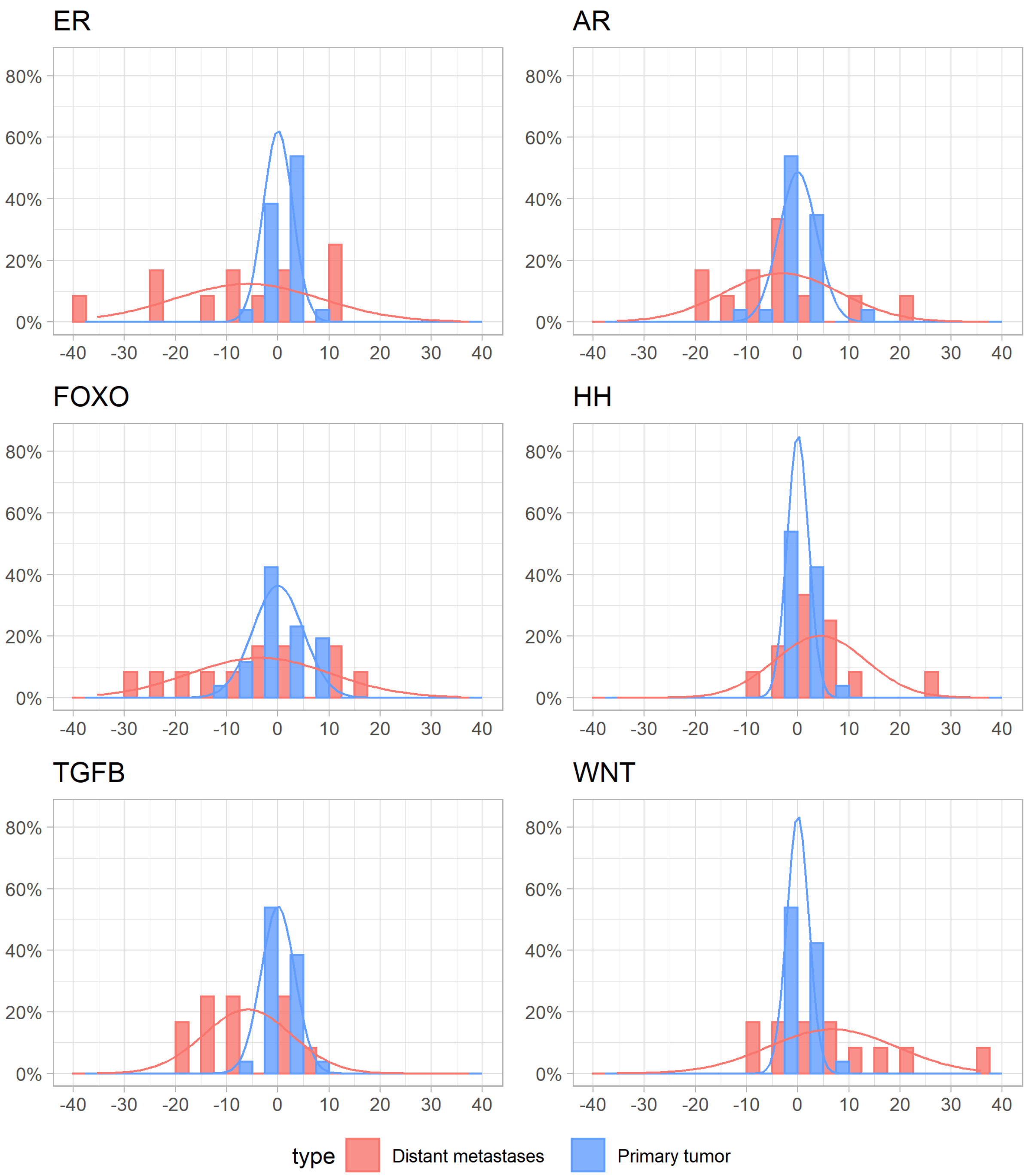


*Figure S-S7: Comparison of percent frequency counts of the metastatic sample specific contributions, $\Delta_{q,meta_k}$ (in red), and of the primary block tissue sample specific contributions, $\Delta_{p,block_j}$ (in blue, same as in Figure S-S6), for the various signaling pathway models. The bars show the percent frequency counts. As a visual guide, solid lines are added representing Gaussian curves with mean and standard deviation equal to the mean and standard deviation values computed for , $\Delta_{q,meta_k}$ and $\Delta_{p,block_j}$, respectively.*

Due to the low sample count, it is difficult to draw conclusions; however, the following observations can be made:

- For all pathways, the spread in LN metastasis specific contribution is generally larger than the spread in primary tumor specific contribution, and the spread in distant metastasis specific contribution is generally larger than the 22% expected solely because there is only a single primary tumor sample available for the distant metastases. This suggests that a metastasis does not necessarily resemble the primary tumor it originates from.
- Besides for the TGFb pathway, compared to LN metastasis, the spread in distant metastasis specific contribution is also larger than the 22% expected solely because there is only a single primary tumor sample available for the distant metastases. This suggests that, with a possible exception of the TGFb pathway, distant metastasis are further away to PT than LN metastasis. (See also correlations in main text.)
- The ER pathway shows an asymmetric distribution for the metastases, with a bias towards lower ER activity in both the LN and the distant metastasis. The few increasing pathway scores shows it is uncommon to gain ER pathway activity in (distant) metastases.
- For the AR pathway, the distribution of the LN metastases appears to be slightly asymmetric (with respect to zero) with a preference for higher AR activity, however, differences are very small. While the distribution of the distant metastases appears to be slightly asymmetric (with respect to zero) with a preference for lower AR activity, however, differences are very small.
- For FOXO activity (PI3K pathway activity interpreted as the inverse of FOXO) the differences are less prominent than for the ER pathway, though FOXO activity seems to be more frequently lost (associated with increased PI3K) in the metastases than gained.
- Distributions for the metastases and primary tumor are similar for the HH pathway, with one clear outlier of a distant metastasis tumor that gained HH activity (located in the ovarium).
- For the TGFB pathway the distribution of the LN and distant metastases appears to be asymmetric with a preference for losing TGFB activity.
- For the WNT pathway the distribution of the LN metastasis appear to be similar to the PT, while distant metastases appears to be asymmetric with a preference for gaining WNT activity), which is inflated by a few outliers that gained WNT activity (metastasis with highest WNT pathway activity score was located in the intestines).
- In general the observations on the lymph node metastases appear to be consistent with, though less pronounced than, observations made on the distant metastases: loss of ER and TGFb pathway activity and of FOXO (indicative of increased PI3K pathway activity).

### 4. Checkerboard model for explaining macro/micro heterogeneity

A simple model for intra-tumor heterogeneity in signaling pathway activity was created. Homogeneous tumor clones are represented as quadrangles and spatially distributed in a checkerboard pattern (Figure S-S7). Assuming the same sample collection procedure is used as in the experimental study, the size of the clones compared to the size of the sampled area is important.

This is used as a parameter in a simulation, where the average “color” of the sampled area is calculated for many iterations, analogous to how the pathway score would average out with an mRNA mixture of two different clones. For example, if a third of the sampled area is white, a value of 1/3 is assigned for that trial. Collected is the standard deviation of many of these trials. Analogous to the experimental setup, after the main sample is collected (macro scale), within the same area, an additional four samples are taken in a quadrant layout (micro scale). Figure S-S7 illustrates the process for two clone sizes.

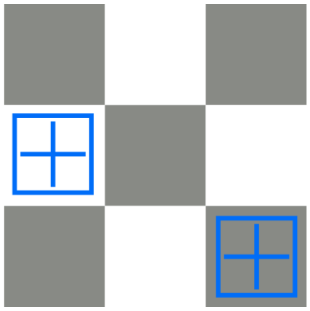

*A. The sampled area represented by a large blue squares simulates a tissue block, and in this example, is smaller than one homogenous cancer clone. There is likely to be heterogeneity between the two simulated samples taken across a tumor. When quadrant samples are analyzed from a single block, they are all from the same cancer clone and expected to show low heterogeneity.*

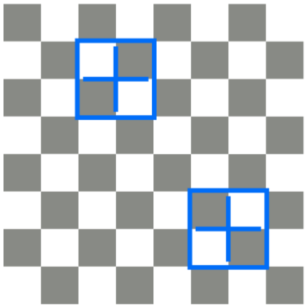

*B. The sampled area represented by a large blue squares simulates a tissue block, and in this example, is bigger than the homogenous clone size, causing the measurements to “average out” with little variation between samples. Smaller quadrants taken from the initial tissue block will show higher heterogeneity, because their size is closer to the clone dimensions.*

*Figure S-S7:* Checkerboard model for cancer clones. *Squares represent cancer cell clones within a tumor in black and white. Analyzing pathway activity in a tissue block results in a pathway score reflecting the average of the pathway activity scores in quadrant samples. If the cancer clones are larger than the tissue block (A), heterogeneity in pathway activity will be found between tissue blocks across the tumor, but not within a tumor block (quadrant samples); in contrast, when cancer clones are smaller than the tissue block (B), heterogeneity in pathway activity will be found within a tissue block.*

The standard deviation of these two different ways of sampling is plotted in Figure S-S8 below. A low ratio (left) indicates the clone size is much bigger compared to the sampling size, and consequently a very high ratio indicates the clone size is very small and fits many times within the sample area.

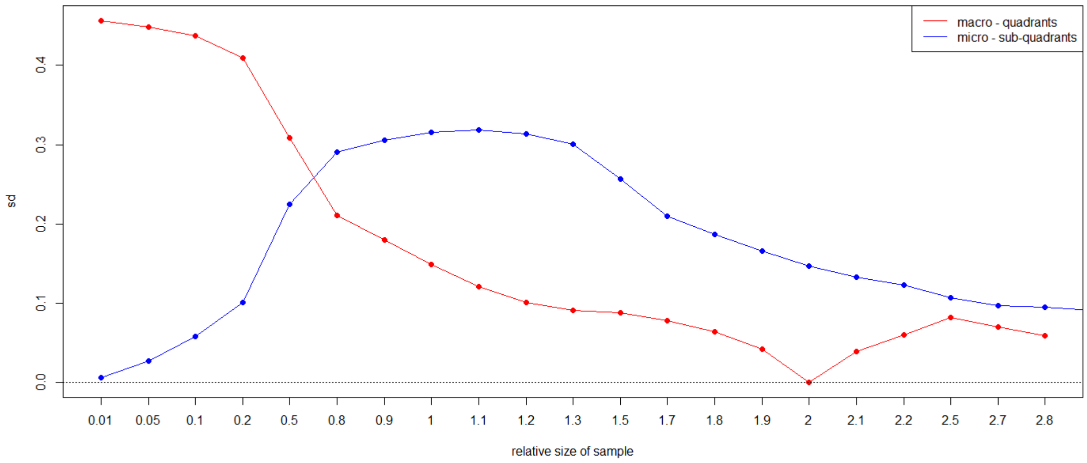


*Figure S-S8: Simulated standard deviation as a function of relative sample size, i.e. the ratio between the sampled area and the clone size.* Macro – quadrants (red): tissue block ; micro – sub-quadrants: quadrants from the tissue block.

From the varying ratios, we can identify a few interesting phenomena, on the premise that the clone size/distribution is even across the tumor:

- In the case of a very low size ratio (sample size smaller than clone size), the micro scale sample variance is much smaller than the macro scale. This is very logical, because if your macro sample is within a large clone, you cannot expect to find any other color within that macro slice. Thus, if the tumor clone is larger than the sampled tissue block, within the tissue block no heterogeneity in pathway activity will be found.
- In the case of a very high size ratio (small clone size, and sample size larger than clone size), the micro scale and macro scale measurements get very similar in size. Again, this is very logical, because the tumor clones are now so small it does not matter if you take micro scale samples or not because the measured average "color" will always be "gray".
- The interesting case is when the size of the tumor clone is comparable to the macro scale sample, i.e. if the tumor clone is similar in dimensions as the scraped area on the slide. In such a case, you can expect much higher micro scale heterogeneity then macro scale heterogeneity. Additionally, if the ratio is known, it could be used to provide an indication of the tumor clone size compared to the scraped area.

#### 5. Choice of software package

There are many statistical packages that enable generation of the model used (Stata, R with lme4 or nlme package, Matlab, SPSS, Minitab). There are two features used beyond the "standard use"; first, the ability to have the lowest level (errors) variance depend on groups (heteroscedasticity), and second, to do inference on (non-linear) functions of the resulting variances. Stata and R can do both.

We show a few snippets of code for these calculations in Stata. To run the micro-macro the model, using cancer subtype T, response value for some pathway `score', and tissue block/quadrant type indicated by sampletype:

```
mixed `score' i.T  || patient: ||block: ||, res(,by(sampletype))reml
```

Here, i.T indicates that type T is a categorical variable. The || part indicates hierarchical random effects. The “res” option specifies that the residual standard deviation has separate values for different groups (in the case either tissueblock samples or quadrant samples).

The same model can be fitted in R using the nlme package function lme (assuming the data is stored in a data frame called data):

```
lme(score  ~ 0 + T, random= list(patient = ~1, block = ~1),
weights=varIdent(form=~1|sampletype) data=data, method="REML")
```
